# Single-cell immune repertoire atlas maps coordinated circulating adaptive immune states in inflammatory bowel disease

**DOI:** 10.64898/2026.09.20.753043

**Authors:** John Gubatan, Jiayu Ye, Christin Lund-Andersen, Jorge Canas, Yuxi Zhou, Theresa Boye, Jacqueline Hoang, Raoul S. Sojwal, Touran Fardeen, Tracy Tran, Peacha Sokzini, Ally Hamin Koh, Marielle Gibson, Yuting Huang, Kathryn Peterson, Sidhartha R. Sinha, Stephan Rogalla, Ole Haagen Nielsen, Michael J. Rosen, Geir Kjetil Sandve

## Abstract

Single-cell studies have defined immune states in inflammatory bowel disease (IBD), but how adaptive receptor histories organize circulating immunity remains unclear. We generated a single-cell transcriptomic atlas of peripheral blood from 249 participants with Crohn’s disease, ulcerative colitis, or non-IBD control status, including 182 with productive TCR and BCR recovery. Expanded TCR clonotypes marked inflammatory-memory and cytotoxic states, while distinct but similar paired TCRs shared inflammatory programs across participants. BCR lineage maturation linked IgA-associated mucosal and plasma B cell programs to somatic mutation and class switching, distinguishing maturation-associated biology from clonal expansion. Helper, regulatory, and cytotoxic T-cell programs covaried with B-cell states, and inferred interactions nominated reciprocal antigen-presentation and helper pathways. Repertoire-based machine learning distinguished diagnosis, inflammation, and contemporaneous six-month treatment-response status. Together, this atlas connects receptor architecture to coordinated systemic immune remodeling, establishes a foundation for repertoire-informed patient stratification, and prioritizes candidate mechanisms of IBD pathogenesis.

**GRAPHICAL ABSTRACT:** 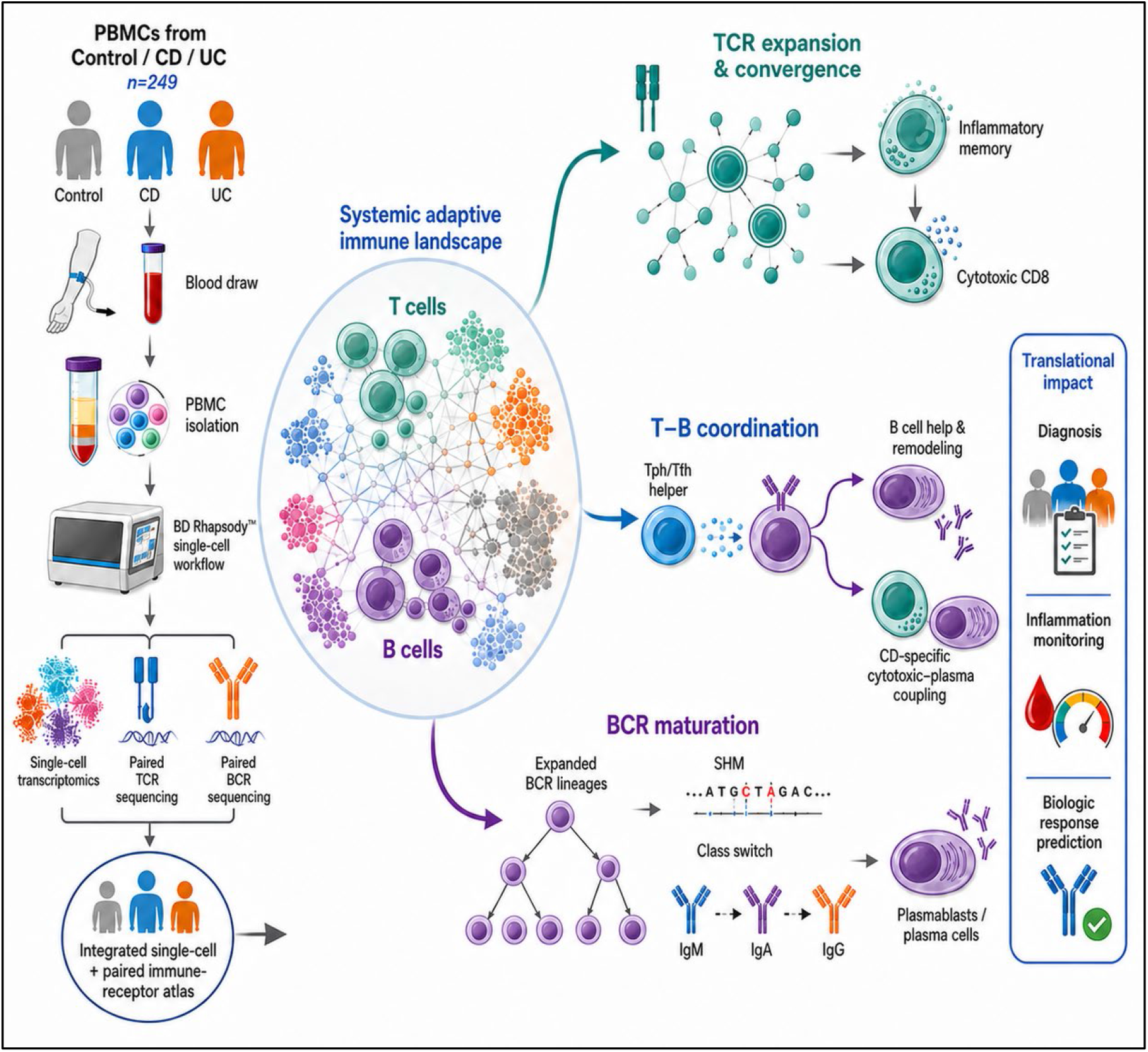

**Highlights:**

- Paired single-cell receptors map circulating adaptive immune states across IBD
- TCR expansion and paired-chain convergence identify inflammatory programs
- BCR lineage maturation links plasma programs to mutation and class switching
- Repertoire features benchmark diagnosis and clinical-state classification

**In Brief:** Gubatan *et al.* present a peripheral blood single-cell transcriptome and paired immune-receptor atlas across IBD and controls. Clone-aware analyses connect TCR expansion and sequence similarity to inflammatory programs, distinguish B-cell phenotype from lineage maturation, and link participant-level T–B covariation to an inferred ligand– receptor interactome. Machine learning models with repertoire features classify diagnosis, inflammatory status, and contemporaneous six-month treatment-response status.

## INTRODUCTION

Inflammatory bowel disease (IBD), comprising Crohn’s disease (CD) and ulcerative colitis (UC), arises from interactions among genetic susceptibility, environmental exposures, the intestinal microbiota, and dysregulated immune responses [1–4]. Its expanding global burden underscores the need to understand the immune processes that sustain disease [5]. The chronic, relapsing course of IBD also raises a question of immune persistence: how previous antigen encounters shape the adaptive immune populations associated with ongoing or recurrent intestinal inflammation. Clinical symptoms, C-reactive protein, fecal calprotectin, and endoscopic assessment provide complementary measures of disease activity, but do not resolve the clonal populations or cellular programs underlying those measurements [6–9]. Connecting prior immune experience to present immune-cell states could therefore provide insight into the organization of persistent inflammation.

Adaptive immune memory provides a biological framework for this connection, through the persistence of antigen-experienced lymphocytes and their capacity to respond to subsequent stimulation. T-cell receptor (TCR) and B-cell receptor (BCR) repertoires capture complementary dimensions of the clonal organization and maturation associated with immune experience [10,11]. Within a participant, exact paired TCR sequences identify clonotypes whose expansion can be related to functional specialization. BCR sequence variation, distance from germline, and immunoglobulin isotype provide additional information about accumulated mutation and maturation. Similarity between nonidentical receptors can nominate potentially related recognition properties, although it does not establish shared antigen specificity [12,13]. Single-cell linkage of receptors to transcriptional profiles connects these features of immune history to current cellular phenotype, allowing clone abundance and receptor architecture to be distinguished from activation, differentiation, and regulatory programs [14]. This distinction is particularly relevant in blood, where antigen-experienced cells coexist with naive and activated populations, and where repertoire measurements alone cannot establish functional immune memory.

Studies of IBD provide evidence that these relationships extend across intestinal and circulating immune compartments. Single-cell analyses of intestinal lesions have identified disease-associated cell states and multicellular programs in CD and UC [15–17]. Repertoire studies have reported altered TCR structure, blood–gut overlap, mutation-associated disease signatures, and shared or antigen-reactive clones [18–25]. Antibody studies likewise connect humoral recognition to intestinal microbes and disease-associated epitopes [26,27]. However, these observations do not yet establish how exact clonal expansion, nonidentical paired-receptor sequence relatedness, and germline-aware BCR maturation relate to circulating T- and B-cell programs within the same participant-level resource. In particular, it remains unclear which receptor–program relationships reflect specialization within comparable cell states, which arise partly from differences in cellular representation, and which recur across participants and independent datasets.

Here, we integrate single-cell transcriptomes and paired immune receptor sequences to examine how clonal expansion, sequence relatedness, and accumulated maturation relate to circulating immune-cell specialization in IBD. We distinguish expansion-associated programs within annotated cell states from changes in cell-state abundance, examine whether nonidentical paired TCRs associate with related programs across participants, and separate BCR distance from germline from diversification among sampled descendants. We then assess how T-cell and B-cell programs covary across individuals and evaluate the information that repertoire sequence features contain about diagnosis and contemporaneous clinical status. This framework connects features of adaptive immune history to present circulating immune states, while distinguishing observed immune organization from mechanisms that may sustain intestinal inflammation.

## RESULTS

### Cohort design anchors the circulating IBD immune-repertoire resource

We profiled 249 participants, including 127 with CD, 87 with UC, and 35 non-IBD controls (Table S1). Median age was 37 years (IQR, 28-49; range, 6-79), and 133 participants (53.4%) were female. Objective intestinal inflammation and biologic exposure were defined prospectively for analysis, enabling diagnosis, inflammatory-state, and contemporaneous post-treatment response-status comparisons. These strata provide clinical context rather than a balanced trial design; endpoint-specific denominators accompany every analysis.

### A clone-aware peripheral-blood atlas maps circulating immune organization

We integrated BD Rhapsody single-cell transcriptomics with paired TCR and BCR sequencing across 249 participants and 10 acquisition series (Figure 1A,B). Transcriptomic profiles were available for the full cohort; 182 participants had productive TCR and BCR calls (control, n=24; CD, n=91; UC, n=67). Productive receptors were linked to cell states and clinical metadata through stable de-identified identifiers. Participants without productive receptor recovery were excluded from receptor analyses rather than encoded as biological zeroes (Figure 1D; Figure S1).

**Figure 1.**
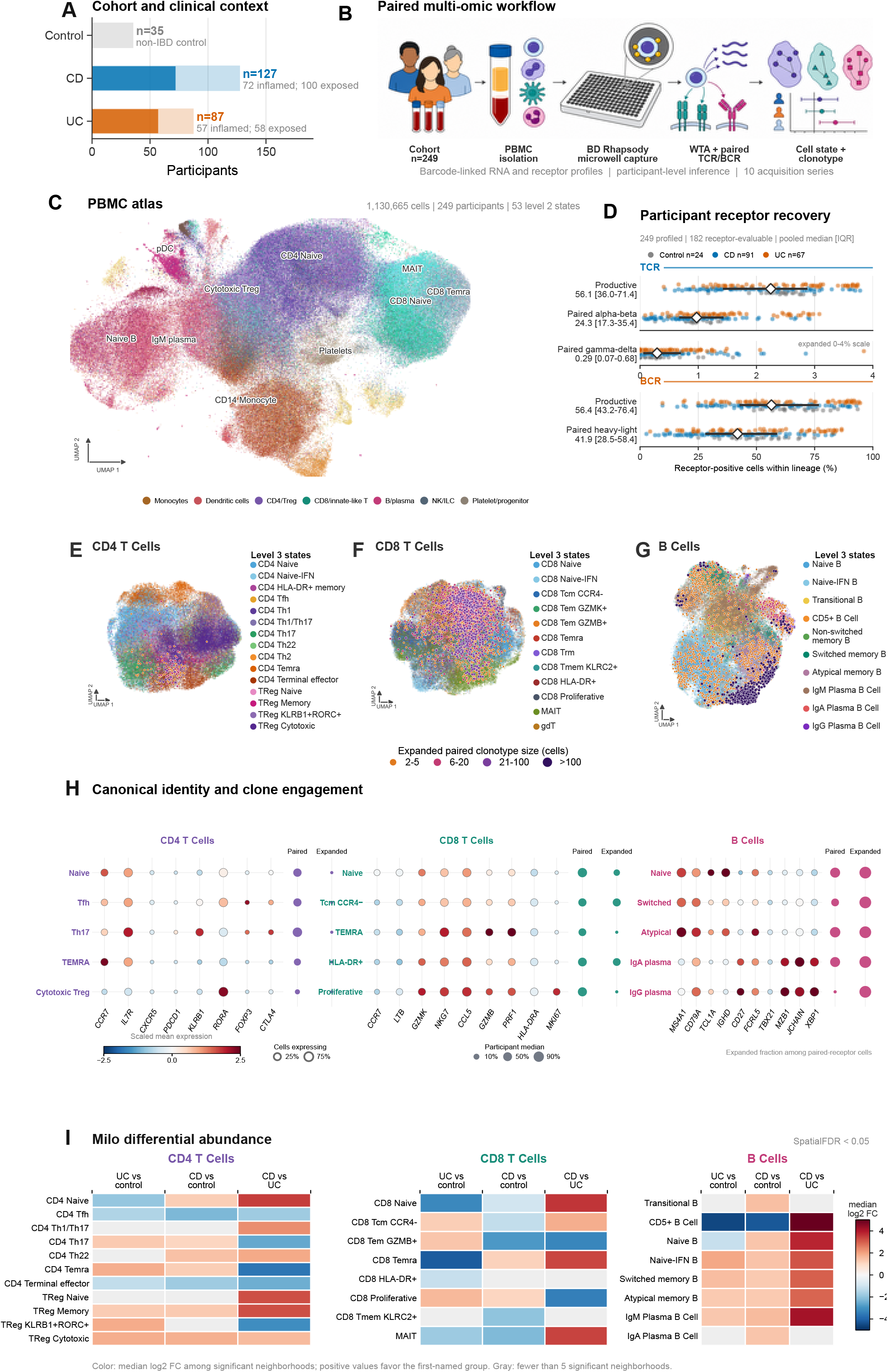
Study design and clone-aware atlas of circulating immune states in IBD. (A) Diagnosis, objective intestinal inflammation, and biologic exposure among 249 participants. Saturated segments indicate inflammation. CD, Crohn’s disease; UC, ulcerative colitis. (B) PBMC transcriptome and paired-receptor sequencing workflow across 10 acquisition series. (C) PBMC UMAP showing 53 level 2 immune states. (D) Participant-level productive receptor recovery (24 controls, 91 CD, and 67 UC). Points represent participants; diamonds and lines indicate medians and interquartile ranges (IQRs). Paired γδ-TCR recovery uses a 0%–4% scale. (E–G) CD4 T-cell (E), CD8 T-cell (F), and B-cell (G) UMAPs showing level 3 states and expanded paired clonotypes. Overlay color and size indicate 2–5, 6–20, 21–100, or >100 cells. Exact paired clonotypes share productive V/J genes and CDR3 amino-acid identities across both receptor chains within a participant; expansion requires ≥2 cells. Downsampling was used only for visualization. (H) Marker expression and receptor engagement across representative states. Dot size indicates the percentage expressing each marker; color indicates scaled mean expression. Receptor columns show participant-median paired-receptor recovery and expanded-cell fractions among paired-receptor cells. (I) Milo neighborhood differential abundance. Color shows the annotation-weighted median log₂ fold change among significant neighborhoods (SpatialFDR <0.05); positive values favor the first-named group. Gray indicates fewer than five significant neighborhoods. These summaries do not test whole-state proportions.

The integrated atlas comprised 1,130,665 quality-controlled cells and 53 level 2 immune states, with higher-resolution annotations for CD4 T cells, CD8 and innate-like T cells, and B cells (Figure 1C,E–G). Marker expression, productive paired-receptor recovery, and expanded-clonotype engagement supported the annotated adaptive states (Figure 1H; Figure S1). Productive recovery and expansion varied by state; missing receptor recovery was not treated as absence of a clone.

Milo differential-abundance analysis identified shared and disease-associated remodeling within circulating adaptive immune states. Relative to controls, neighborhoods annotated as cytotoxic Tregs were consistently enriched in both CD and UC, whereas MAIT and CD5⁺ B-cell neighborhoods were predominantly depleted. Naive– IFN B-cell and IgM-plasma neighborhoods also generally favored IBD. Direct comparison of the two diseases revealed enrichment of CD4 Th1/Th17 neighborhoods in CD, whereas KLRB1⁺RORC⁺ Treg and GZMB⁺ CD8 effector-memory neighborhoods predominantly favored UC. In contrast, conventional CD4 Th17 neighborhoods exhibited substantial bidirectional changes, indicating heterogeneity within this annotation rather than uniform population expansion. Together, these findings identify shared and divergent local immune-state changes in IBD and provide a cellular context for subsequent receptor-resolved analyses. Because Milo tests overlapping transcriptional neighborhoods, these results should not be interpreted as direct measurements of whole-state proportions (Figure 1I; Tables S2–S4). The atlas therefore provides the cellular context needed to ask whether IBD-associated repertoire features reflect changes in state representation, within-state transcriptional programs, or both.

### Exact TCR clonotypes define a circulating inflammatory-memory and cytotoxic axis

We first characterized global TCR repertoire architecture across 91 participants with CD, 67 with UC, and 24 controls. Shannon diversity summarized the richness and evenness of recovered receptor sequences, whereas clonality quantified their concentration within dominant sequences. Median clonality was higher in UC and CD than in controls (0.124, 0.067, and 0.007, respectively), accompanied by lower median Shannon diversity (6.09, 6.28, and 6.76; Figure 2A). Additional diversity measures supported greater inequality in sequence representation: median Gini coefficients were 0.592 in UC, 0.493 in CD, and 0.059 in controls, while median inverse Simpson diversity was 150.6, 310.1, and 598.5, respectively. Median sequence richness, however, was higher in UC and comparable between CD and controls (1,128, 920, and 916.5, respectively), indicating that reduced diversity did not reflect a uniform loss of recovered sequence richness (Figure S2A). CDR3 amino-acid–aggregated repertoire counts further revealed differences in clone-size composition. Controls were predominantly represented by singleton counts, whereas medium (6–20) and large (21–100) count bins contributed more prominently to IBD repertoires; the hyperexpanded (>100) bin had its highest median contribution in UC (Figure S2B). Analysis of exact paired αβ clonotypes additionally showed that expanded and non-expanded compartments differed in cellular composition across all three diagnostic groups (Figure S2C). CD4 naive cells constituted approximately 39–40% of the mean non-expanded compartment, whereas expanded compartments were predominantly composed of CD8 states. CCR4-negative CD8 central memory cells accounted for mean proportions of 40.5% in UC, 41.9% in CD, and 55.8% in controls within the expanded compartment, compared with 6.2%, 4.7%, and 4.6%, respectively, within the non-expanded compartment. CD8 HLA-DR+ cells and cells annotated as CD8 naive also made substantial contributions to expanded compartments. CD8 Temra cells contributed approximately 7.5–7.7% of the expanded compartment in IBD. These compositional findings show that expanded paired clonotypes occupied several cellular states and that CD8 predominance was also present in controls. Together, the results distinguish repertoire concentration, cellular expansion burden, and the cellular identity of expanded clones, motivating subsequent analyses of clone-associated transcriptional programs within matched T-cell states.

**Figure 2.**
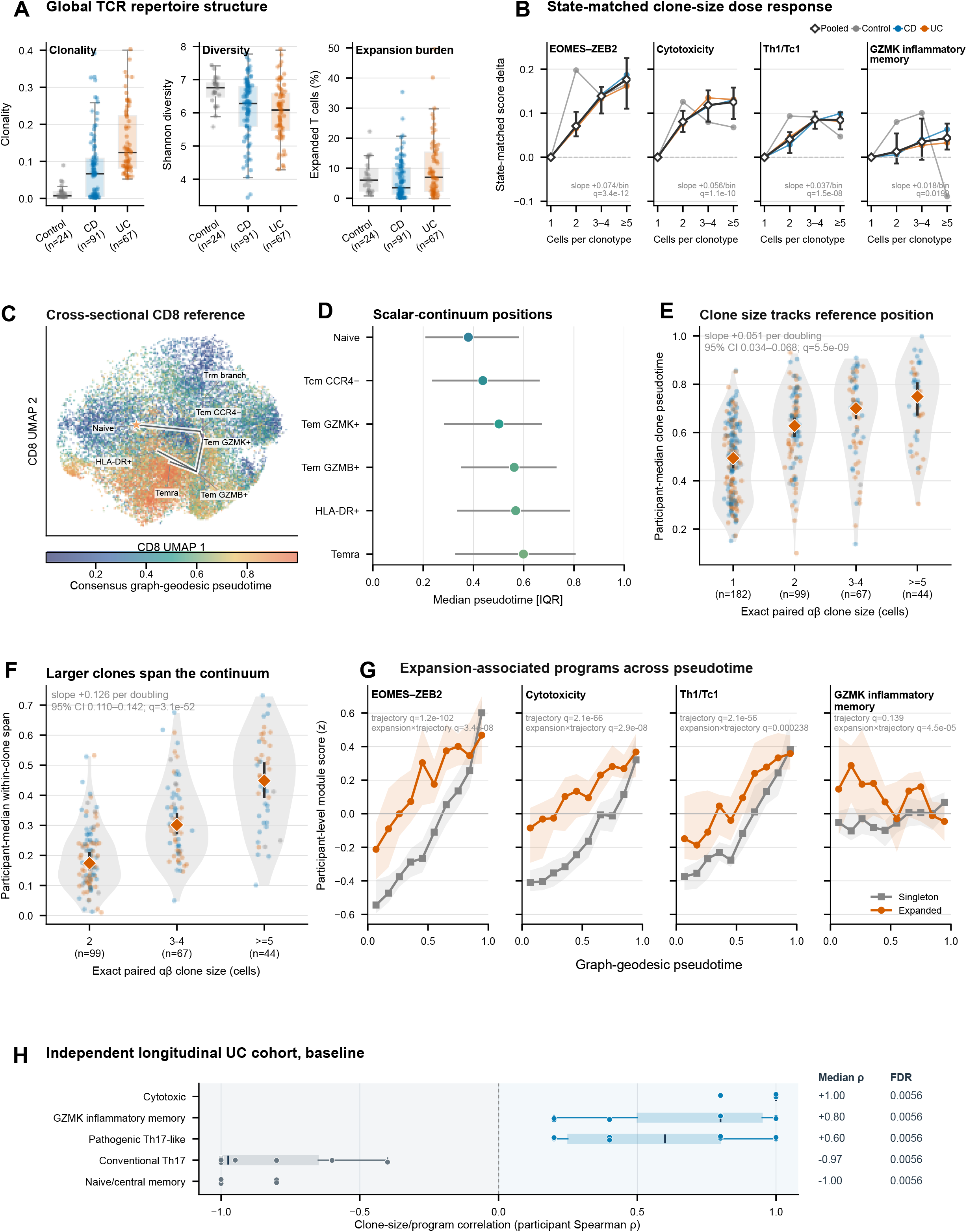
TCR expansion associates with inflammatory-memory and cytotoxic states. (A) Participant-level CDR3 amino-acid repertoire clonality and Shannon diversity, and expanded-cell fractions under the exact productive TCRβ definition. Boxes show medians and IQRs. (B) State-matched program-score differences relative to singleton cells across exact paired αβ clone-size bins. States and CD4/CD8 compartments were equally weighted within participants. Intervals show participant-bootstrap 95% confidence intervals (CIs); annotations indicate ordered-trend FDRs. (C,D) Conventional-CD8 graph-geodesic pseudotime (C) and state medians with IQRs (D). (E,F) Clone-median pseudotime (E) and within-clone 90th–10th percentile span (F), summarized by participant. Diamonds and bars show pooled medians and participant-bootstrap 95% CIs. Slopes estimate within-participant effects per clone-size doubling. Regression samples included 109 and 71 participants, respectively, and differ from displayed bin counts. (G) Participant-balanced program profiles for singleton and expanded paired αβ clonotypes. Shading indicates participant-bootstrap 95% CIs; interaction FDRs derive from participant-blocked spline models. (H) Baseline validation in an independent UC cohort [28]. Points show participant-level Spearman correlations between four ordered clone-size bins and mean program score (n = 10); boxes show medians and IQRs. Two-sided signed-rank tests against zero were FDR adjusted across the external clone-size/program family. External analyses were not state matched. Pseudotime represents transcriptional ordering, not observed temporal progression. See also Figures S2A–C and S3.

We asked whether clone size was associated with inflammatory and cytotoxic programming beyond differences in broad T-cell identity. Comparing exact paired αβ-TCR clonotypes within the same participant and annotated T-cell state revealed progressively higher EOMES–ZEB2, cytotoxicity, Th1/Tc1, and GZMK inflammatory-memory scores across singleton, 2-cell, 3–4-cell, and ≥5-cell clones (ordered-trend FDR = 3.40 × 10⁻¹², 1.12 × 10⁻¹⁰, 1.48 × 10⁻⁸, and 0.0199, respectively; Figure 2B). Thus, larger clones were distinguished not only by their abundance but also by inflammatory-memory and effector features within otherwise similar cell states.

An independent UC cohort [28] supported this relationship while resolving an important distinction within Th17-associated biology. At baseline, larger clones were associated with cytotoxic, GZMK inflammatory-memory, and pathogenic-Th17 programs, but inversely associated with conventional-Th17 and naive/central-memory programs (median participant-level ρ = 1.00, 0.80, 0.60, −0.97, and −1.00, respectively; all FDR = 0.0056; n = 10; Figure 2H). These external analyses were not state matched and therefore provide complementary evidence linking clone size to inflammatory effector features, rather than demonstrating generalized expansion of conventional Th17 cells. Baseline disease-control differences and week-6 changes did not survive FDR correction, and response associations could not be evaluated without participant-linked response labels (Figure S3).

Within conventional CD8 T cells, transcriptional ordering placed larger paired αβ clonotypes farther along an early-memory-to-cytotoxic continuum, with a separate tissue-resident-memory-like branch (Figure 2C,D). Clone-median pseudotime increased by 0.051 per clone-size doubling (95% CI, 0.034–0.068; FDR = 5.49 × 10⁻⁹; Figure 2E). Expanded clones also spanned broader transcriptional positions: their 90th–10th percentile pseudotime range increased by 0.126 per doubling (95% CI, 0.110–0.142; FDR = 3.05 × 10⁻⁵²; Figure 2F). Larger clones therefore combined a later effector-associated position with greater within-clone state diversity, rather than occupying a single terminal phenotype. Expansion-associated differences in all four inflammatory and effector programs varied along pseudotime (interaction FDR ≤ 2.38 × 10⁻⁴; Figure 2G). Together, these findings connect clonal abundance to both inflammatory specialization and transcriptional heterogeneity, providing a framework for understanding how circulating adaptive immunity is remodeled in IBD. Cross-sectional pseudotime, however, does not establish temporal differentiation or show that expansion causes these programs.

### Paired-TCR sequence convergence recovers expansion-linked inflammatory programs

The expansion analyses linked clone size to inflammatory specialization within matched cellular states. We next asked whether related transcriptional programs also occurred across participants carrying similar but nonidentical paired TCRs, extending the analysis beyond the abundance of individual clones. A sequence-similarity graph connected 43,742 participant-specific paired αβ clonotypes across eight acquisition series, identifying 5,548 cross-participant links involving 181 participants (Figure 3A). Similarity incorporated both chains’ CDR3 amino-acid features and V/J-gene usage, while excluding identical receptor pairs. This design tested whether shared immune states extended beyond recurrence of the same clonotype.

**Figure 3.**
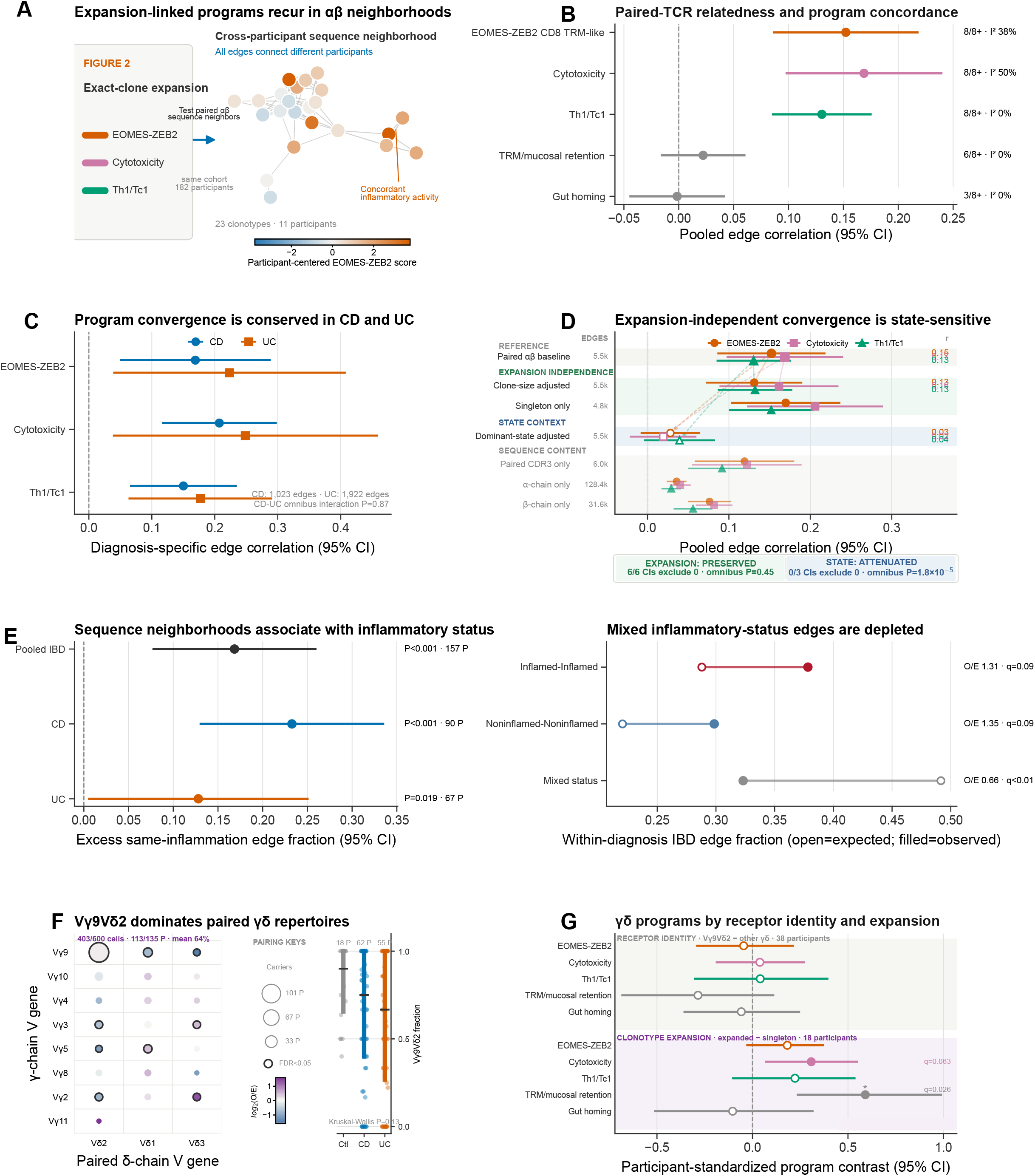
Paired-TCR sequence similarity links inflammatory programs across participants. (A) Cross-participant sequence neighborhoods comprising 43,742 paired αβ clonotypes from 182 receptor-evaluable participants. The displayed network is the acquisition-series S3 component containing the most participants. Edges connect non-identical receptors across participants; width indicates similarity and node color indicates participant-centered EOMES–ZEB2 activity. (B) Random-effects correlations between sequence neighbors for five programs. Points and bars show pooled estimates and participant-jackknife 95% CIs (5,548 edges; 181 participants). (C) CD- and UC-restricted correlations for three primary programs; the omnibus diagnosis interaction was nonsignificant (P = 0.87). (D) Sensitivity to clone size, singleton restriction, dominant cell state, and receptor-chain definition. Filled symbols indicate CIs excluding zero. Joint attenuation tests compare adjusted and baseline correlations. (E) Inflammatory-status organization among 3,246 within-diagnosis IBD edges from 157 participants. Null expectations derive from 10,000 within-diagnosis participant-label permutations. (F) Participant-normalized γδ V-gene pairing across 135 participants. Bubble area indicates participant prevalence, color indicates log₂ observed-to-expected pairing, and outlines indicate FDR <0.05. (G) State-matched program contrasts for TRGV9–TRDV2 versus other γδ receptors (n = 38 participants) and expanded versus singleton γδ clonotypes (n = 18). Points and bars show mean participant-level contrasts and bootstrap 95% CIs. Sequence graphs were constructed within acquisition series; γδ contrasts did not include an additional batch covariate. Sequence similarity does not establish shared antigen specificity. See also Figure S2D.

Sequence-related clonotypes showed concordant EOMES–ZEB2, cytotoxicity, and Th1/Tc1 programs across participants (pooled correlations = 0.152, 0.169, and 0.130; Figure 3B). Concordance remained positive within diagnostic groups, without evidence of an overall CD–UC difference (P = 0.87; Figure 3C), suggesting a receptor-associated inflammatory organization not confined to either disease. Importantly, concordance persisted after clone-size adjustment and among singleton clonotypes. It therefore could not be explained solely by the preferential expansion of similar receptors. By contrast, adjustment for dominant cell state attenuated concordance (joint P = 1.85 × 10⁻⁵), whereas clone-size adjustment did not (P = 0.45; Figure 3D). These findings associate paired-receptor relatedness with shared inflammatory programs across participants, including among singleton clonotypes, while the attenuation after cell-state adjustment places part of this relationship within shared cellular identities rather than a demonstrated receptor-intrinsic effect.

Sequence similarity also tracked current inflammatory context. Among 3,246 within-diagnosis links from 157 participants with IBD, connections between participants sharing inflammatory status exceeded the permuted expectation by 16.8 percentage points (95% CI, 7.8–25.9; P = 0.0006). This organization was detectable separately in CD and UC (P = 0.0002 and 0.019; Figure 3E). Thus, receptor neighborhoods captured an association with contemporaneous inflammation beyond diagnostic grouping. External comparison with five published GLIPH2 specificity motifs [29,30] provided context but not validation of these neighborhoods: two motifs were enriched in the published European cohort, whereas none survived FDR correction internally (Figure S2D).

The γδ compartment exhibited a different pattern, dominated by TRGV9–TRDV2 receptors across diagnoses (403 of 600 cells; 113 of 135 participants; diagnosis comparison P = 0.13; Figure 3F). Within matched γδ states, expanded clonotypes showed higher mucosal-retention program activity (effect = 0.591; FDR = 0.026; n = 18), whereas receptor-identity contrasts were not FDR significant (Figure 3G). Together, these analyses distinguish cross-participant αβ sequence convergence on inflammatory states from a broadly shared γδ receptor architecture with expansion-associated functional variation. They nominate receptor-linked features of systemic IBD inflammation without demonstrating shared antigen specificity, intestinal origin, or tissue trafficking.

### BCR clonal focusing marks branching antibody-secreting states

Circulating BCR repertoires in IBD were more concentrated within dominant receptor sequences, accompanied by selective redistribution of expanded cells across B-cell states. Both clonality and clonal-expansion indices were higher in CD and UC than in controls (clonality FDR = 0.0114 and 9.33 × 10⁻⁹; expansion-index FDR = 2.59 × 10⁻⁶ and 7.91 × 10⁻¹³, respectively; Figure 4A). These related repertoire-concentration measures were complemented by paired-chain analyses showing enrichment of selected IgA- and IgM-plasma, memory, transitional, and CD5⁺ states among expanded cells, with depletion of naive and atypical-memory states (Figure 4B; Figure S4). Thus, repertoire focusing was associated with specific cellular phenotypes rather than uniform expansion across the B-cell compartment.

**Figure 4.**
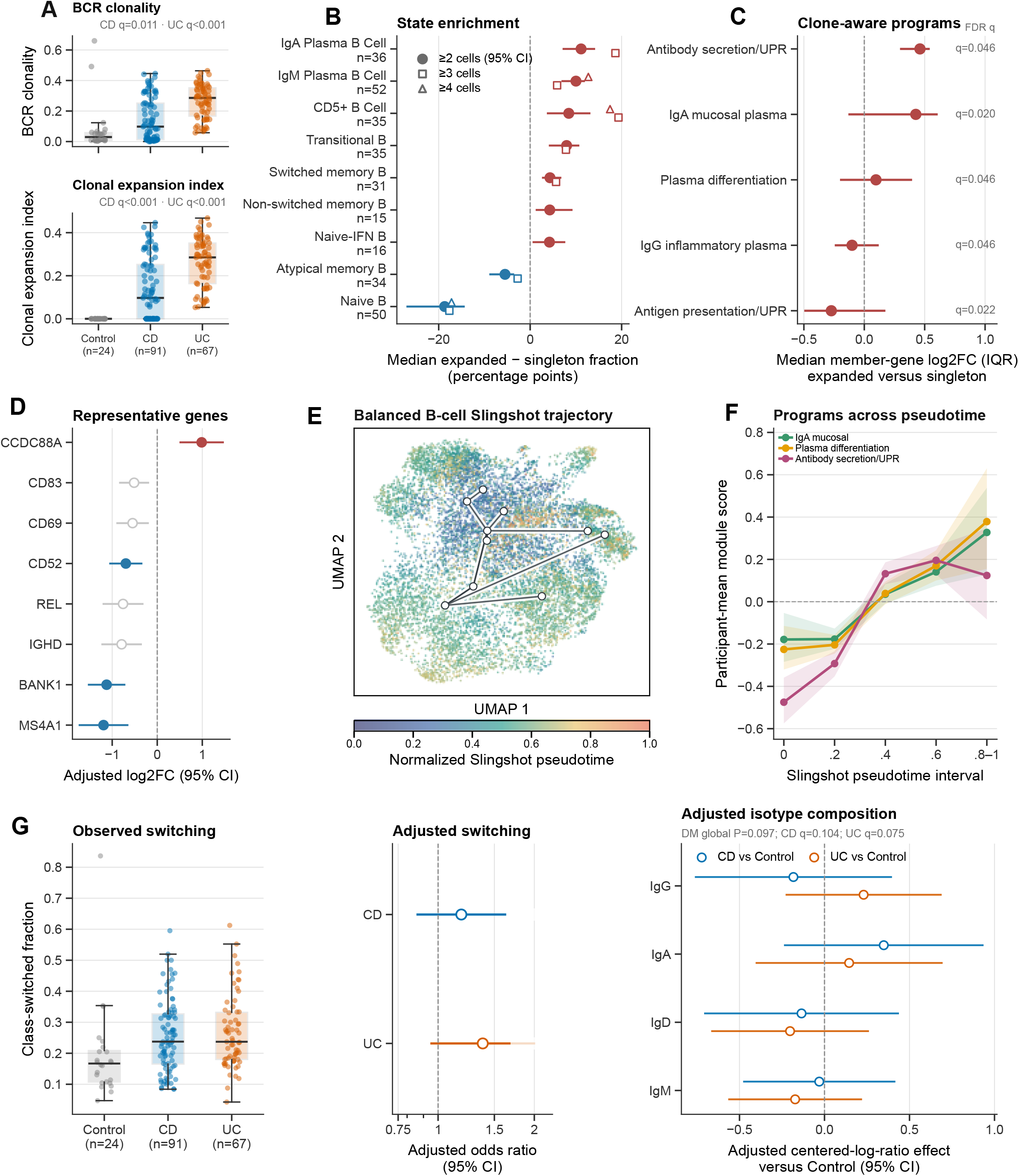
BCR repertoire focusing associates with distinct plasma-cell programs. (A) Participant-level BCR clonality and expansion index. Boxes show medians and IQRs; disease-control comparisons used two-sided rank-sum tests with FDR correction within metric. (B) Expanded-minus-singleton state fractions, defined using heavy-chain V/J genes and paired heavy/light CDR3 identities. Circles and bars show ≥2-cell effects and participant-bootstrap 95% CIs; squares and triangles indicate ≥3- and ≥4-cell thresholds. (C,D) Participant-paired, state-matched pseudobulk comparisons using heavy-chain-defined expansion (n = 38 participants), adjusted for RNA complexity and mitochondrial fraction. In (C), points and lines show median and IQR member-gene log₂ fold changes; q values are competitive cameraPR FDRs, not tests of the displayed medians. In (D), points and bars show representative gene effects and 95% CIs; filled points indicate global FDR <0.05. (E) Slingshot trajectory of 14,868 participant/state-balanced B cells, fitted in 10-dimensional scVI space with a transitional root and four terminal-state constraints. (F) Participant-balanced standardized programs across branch-weighted pseudotime; shading indicates participant-bootstrap 95% CIs. (G) Observed switched fractions, adjusted switching odds ratios, and centered-log-ratio isotype coefficients. Models adjusted for age, sex, and BCR depth. Switching contrasts used beta-binomial models, with displayed P values adjusted by the Holm method; overall isotype composition used a Dirichlet–multinomial test. Pseudotime and isotype occupancy do not establish temporal differentiation or switching direction. See also Figures S4–S6.

To distinguish transcriptional differences within states from changes in their relative abundance, we compared expanded and nonexpanded B cells within participants and matched cell states. This complementary pseudobulk analysis included 38 eligible participants and defined expansion by heavy-chain identity rather than exact heavy–light pairing. Five programs passed competitive gene-set FDR correction. Antibody-secretion/UPR, IgA-mucosal, and plasma-differentiation programs had positive median member-gene effects, whereas IgG-inflammatory plasma and plasma-cell antigen-presentation/UPR programs had negative median effects (Figure 4C). Expansion was therefore associated with a selective combination of antibody-producing-cell features, not a coordinated increase in every plasma-cell or stress-response program. Increased CCDC88A and decreased MS4A1, BANK1, and CD52 further distinguished expanded cells (Figure 4D). The opposing program-level summaries and heterogeneous gene effects argue against treating secretory differentiation as a single, uniformly activated process.

A branching transcriptional trajectory placed these phenotypes within a broader landscape of B-cell specialization. Using a participant- and state-balanced reference of 14,868 cells, Slingshot [31] organized cells from a transitional-B-cell root toward atypical-memory and IgM-, IgA-, and IgG-plasma endpoints (Figure 4E). IgA-mucosal, plasma-differentiation, and antibody-secretion/UPR programs followed distinct profiles along this ordering (Figure 4F), separating mucosal-associated identity from antibody-producing-cell and protein-handling machinery. These profiles describe transcriptional specialization; they do not measure antibody secretion or directly establish temporal differentiation.

Repertoire focusing was not accompanied by a clearly independent diagnosis-level increase in class switching. Although observed class-switched fractions were higher in CD and UC than in controls (Holm-adjusted P = 0.00266 and 0.00114), these differences were not significant after adjustment for age, sex, and BCR depth (CD odds ratio = 1.18, 95% CI 0.86–1.62; UC odds ratio = 1.38, 95% CI 0.95–2.00; adjusted P = 0.298 and 0.176). Overall isotype composition also lacked a significant diagnosis effect (P = 0.0969; Figure 4G). Together, these findings link systemic BCR remodeling in IBD to selective cellular and transcriptional specialization rather than a generalized increase in switching activity, motivating lineage-resolved analyses that distinguish clonal abundance from receptor maturation.

### BCR lineages separate accumulated maturation from observed-only diversification

The B-cell expansion analyses identified a selective combination of cellular and transcriptional features rather than uniform activation of plasma-cell programs. We next examined accumulated receptor maturation, asking whether germline-relative heavy-chain somatic hypermutation (SHM) and lineage architecture provided information distinct from the phenotypes associated with clonal expansion. Regional SHM was higher in IBD, with preferential targeting of complementarity-determining regions (CDRs) relative to framework regions (Figure 5A,D). Isotype-matched comparisons localized the clearest increase to IgM: the UC–control difference survived FDR correction, whereas the CD– control difference did not (FDR = 0.0054 and 0.0515, respectively; Figure 5B). Disease-associated heavy-chain SHM differences were also observed across unique-sequence, paired-clonotype, and cell-weighted summaries (Figure 5C). Thus, the maturation-associated signal was not confined to class-switched receptors or a single sequence-weighting definition.

**Figure 5.**
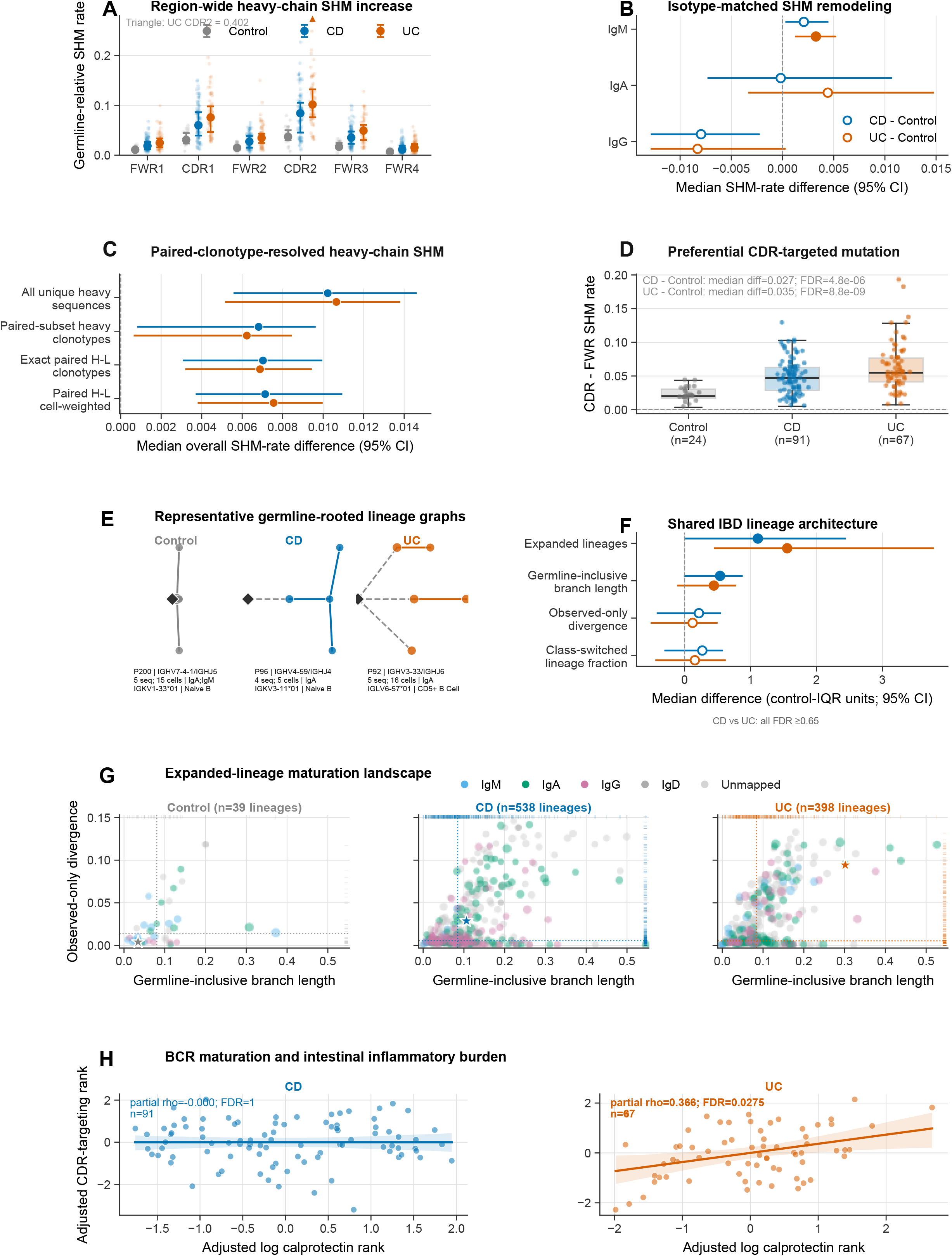
BCR lineages distinguish accumulated mutation from sampled diversification. (A) Heavy-chain somatic hypermutation (SHM) across framework regions (FWRs) and complementarity-determining regions (CDRs) in 24 controls, 91 participants with CD, and 67 with UC. Large points and bars show medians and IQRs; the triangle marks one UC CDR2 value above the plotted range. (B) Isotype-matched disease-control SHM differences. Points and bars show median differences and participant-bootstrap 95% CIs; filled symbols indicate FDR <0.05. (C) SHM differences across unique-sequence, light-chain-linked heavy-clonotype, exact heavy–light, and cell-weighted definitions. Measurements represent heavy-chain SHM, not combined heavy- and light-chain mutation. (D) CDR targeting, defined as mean CDR1/CDR2 SHM minus mean FWR1– FWR4 SHM. (E) Representative germline-rooted lineages selected using prespecified criteria: class switched, paired-light-chain linked, and containing 4–10 observed heavy-chain alignments. Node size indicates abundance; diamonds mark germlines. Dashed and solid edges connect germline and observed sequences, respectively. (F) Control-IQR-standardized disease effects on lineage burden, germline-inclusive branch length, observed-only divergence, and class-switched lineage fraction. Filled symbols indicate FDR <0.05. (G) Expanded-lineage maturation landscape. Point size indicates linked-cell abundance, color indicates dominant isotype, and stars identify lineages in (E); dotted lines mark diagnosis-specific medians. (H) Association between CDR targeting and fecal calprotectin after rank residualization for lineage depth, biologic exposure, and acquisition series. Acquisition series was substantially confounded with diagnosis. Lineage architecture does not establish mutation rate or switching direction. See also Figures S5, S6, and S9.

Germline-rooted lineage reconstruction separated accumulated distance from germline from diversification among the sequences recovered at sampling (Figure 5E–G). Both CD and UC had greater expanded-lineage burden and germline-inclusive branch length than controls (FDR <0.05), but neither observed-only divergence nor the class-switched lineage fraction differed significantly. Representative lineage graphs and the expanded-lineage landscape illustrate these distinct dimensions of receptor history. Greater germline-inclusive branch length without greater observed-only divergence is consistent with more extensively mutated circulating receptors, rather than evidence of an increased current mutation rate. Threshold and lineage-definition sensitivity analyses are provided in Figure S5.

Lineage-linked transcriptional analyses connected this maturation history to selected B-cell programs (Figure S6A). After excluding immunoglobulin constant-region genes and accounting for participant, annotated cell state, lineage abundance, scored-cell depth, and pseudotime, IgA-mucosal and plasma-differentiation activity remained positively associated with SHM/germline distance (standardized effects per program SD = 0.058 and 0.051; FDR = 9.74 × 10⁻²⁰ and 2.69 × 10⁻¹⁶, respectively). Both programs were also associated with class-switching probability, but not with every measure of lineage complexity. These findings distinguish maturation-associated plasma specialization from a generalized increase in sampled diversification or cross-state occupancy.

The relationship with clinical inflammation was selective. CDR targeting correlated with fecal calprotectin in UC after adjustment for lineage depth, biologic exposure, and acquisition series (partial ρ = 0.366; FDR = 0.0275; n = 67), whereas no corresponding association was detected in CD (Figure 5H). Binary objective-inflammation contrasts did not survive FDR correction (Figure S6B). Together, the lineage and transcriptional analyses associate accumulated receptor maturation with selected plasma-cell programs without a corresponding significant increase in diversification among sampled descendants (Figure 5; Figure S6A). This distinction separates the mutation history represented in circulating receptors from evidence of ongoing diversification and clarifies why maturation-associated specialization need not parallel diagnosis-level differences in class switching. The UC calprotectin association further links one aspect of this maturation history to intestinal inflammatory burden. Diagnosis–acquisition-series confounding and the absence of external SHM validation limit disease-level interpretation. Lineage architecture does not establish antigen specificity, mutation rate, or temporal switching direction (Figures S6 and S9).

### Helper and regulatory programs link circulating T-cell activity to humoral specialization

Having related receptor features to cellular specialization within each compartment, we next asked how T-cell and B-cell programs were organized across the same individuals. Participant-level models tested which helper and regulatory programs retained associations with selected B-cell states after accounting for their overlapping information and measured covariates. Participant-level analyses linked TCR and BCR repertoire structure with selected transcriptional features (Figure 6A; Figure S7A–D). Within pooled IBD, pathogenic-Th17, conventional-Th17, suppressive-Treg, and Tph/Tfh-help programs each correlated positively with five B-cell programs (partial ρ = 0.233– 0.590; all 20 correlations FDR ≤ 0.00081; Figure 6B). These associations persisted after adjustment for demographic, clinical, lineage-depth, and acquisition-series covariates. The analysis included 207 participants with evaluable transcriptomes, a larger population than the receptor-evaluable cohort because productive paired-receptor recovery was not required.

**Figure 6.**
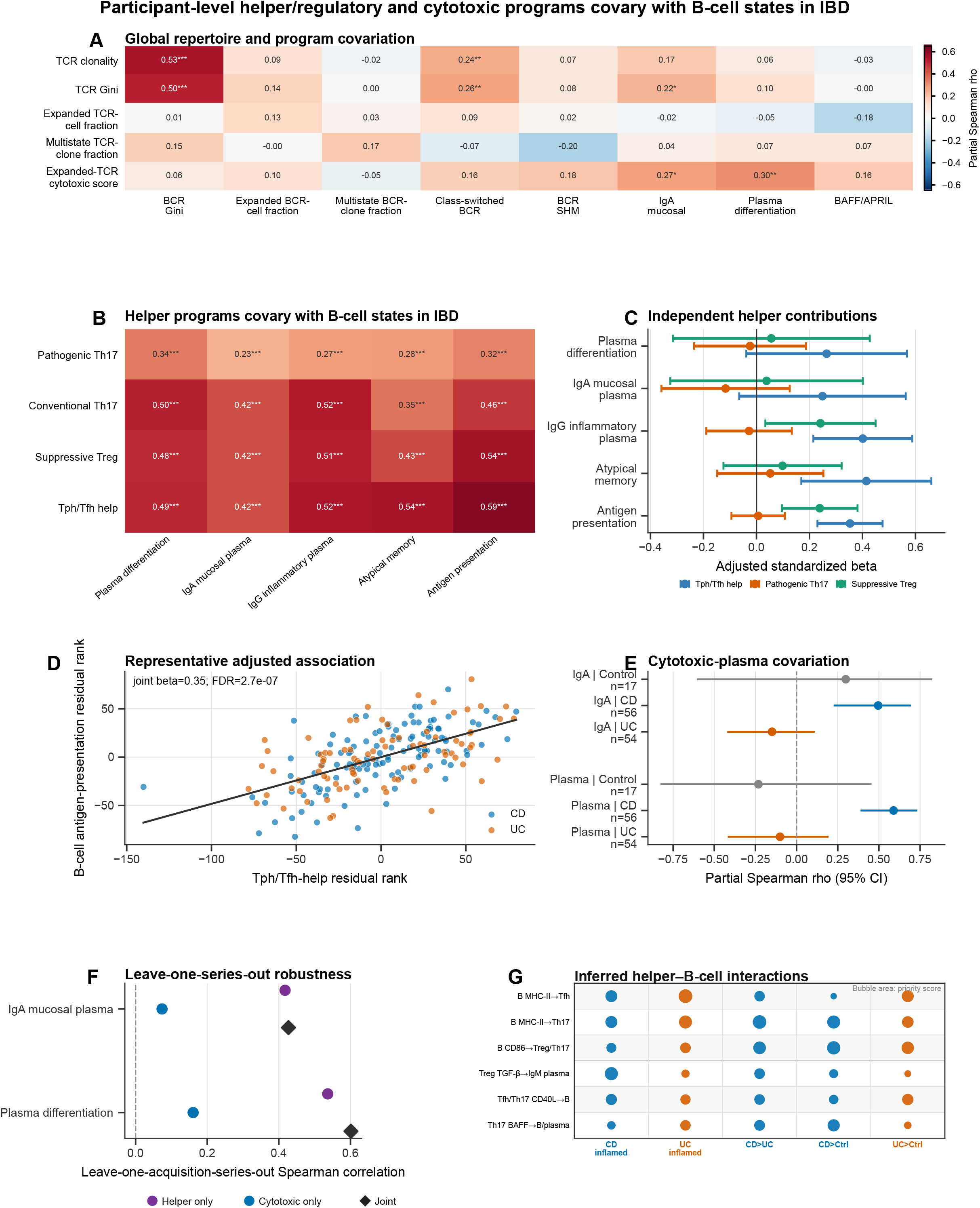
T–B coordination and inferred reciprocal communication in IBD. (A) Participant-level partial Spearman correlations between TCR and BCR features, adjusted for diagnosis, age, sex, and receptor depth. Asterisks indicate FDR <0.05 (*), <0.01 (**), or <0.001 (***). (B) Helper/regulatory–B-cell program correlations in transcriptome-evaluable IBD (n = 207), adjusted for diagnosis, demographics, lineage depth, inflammation, biologic-response status, and acquisition series. Significance used 5,000 series-blocked permutations with FDR correction. (C) Independent helper/regulatory contributions in joint models. Points and bars show standardized coefficients and robust 95% CIs; FDR correction included 15 terms. (D) Adjusted Tph/Tfh-help–B-cell-antigen-presentation association. Axes show rank residuals; the line is a descriptive fit. (E) Diagnosis-stratified correlations between expanded-TCR cytotoxicity and IgA-mucosal or plasma-differentiation programs, with participant-bootstrap 95% CIs. (F) Leave-one-acquisition-series-out evaluation of helper-only, cytotoxic-only, and joint models. Values correlate pooled held-out predictions with observed scores. Joint-minus-helper improvements were not permutation significant. (G) MultiNicheNet-inferred T–B interactions. Rows summarize B-cell MHC-II toward Tfh or Th17, B-cell CD86 toward Treg/Th17, Treg TGF-β toward IgM plasma, Tfh/Th17 CD40L toward B cells, and Th17 BAFF toward B/plasma cells. Columns show CD or UC inflammation contrasts, shared-series CD versus UC, and S6-restricted disease-control contrasts. Each bubble represents the highest-priority eligible pair within its row and context; area encodes priority and color denotes CD or UC. Exact interaction identities and scores are provided in Source Data. Correlations and inferred interactions do not establish direct signaling, antigen specificity, or causality. See also Figures S7 and S8.

Despite this broad covariation, helper programs differed in the information they contributed independently. When modeled together, Tph/Tfh help remained associated with IgG-inflammatory plasma-cell activity, atypical-memory activity, and B-cell antigen presentation (standardized β = 0.401, 0.415, and 0.353; FDR = 0.00019, 0.0039, and 2.70 × 10⁻⁷, respectively; Figure 6C,D). Suppressive-Treg activity also remained associated with antigen presentation (β = 0.239; FDR = 0.0039), whereas pathogenic-Th17 associations were no longer significant. Thus, the shared inflammatory context did not make all T-cell programs interchangeable: Tph/Tfh-associated activity retained a distinct relationship with humoral effector and antigen-presenting states, alongside a regulatory component. The loss of independent pathogenic-Th17 significance indicates overlapping explanatory information, not exclusion of Th17 biology.

This coordination across participants was not accompanied by evidence of preferential clonal sharing between Th17 and Treg states. Only eight exact paired αβ clonotypes contained both states, compared with 14.57 expected on average under within-participant state-label permutation (P = 0.0264; Figure S8A,B). Clone-size and state-matched expansion analyses of the four focused Th17/Treg programs did not survive FDR correction (Figure S8C,D). These findings distinguish coordinated activity across cellular populations from their occupation of the same clonotypes. They neither exclude Th17–Treg plasticity nor establish that shared clonal ancestry underlies helper– B-cell covariation.

A complementary cytotoxic–humoral relationship was strongest within CD. Cytotoxic activity among expanded TCR clonotypes correlated with IgA-mucosal and plasma-differentiation programs (partial ρ = 0.50 and 0.59; n = 56; Figure 6E), connecting an effector T-cell feature to antibody-producing-cell specialization. However, helper activity accounted for much of the reproducible cross-participant information. In leave-one-acquisition-series-out evaluation, helper-only models retained correlations of 0.417 and 0.536 with the two B-cell programs; adding cytotoxicity increased these to 0.427 and 0.601, without significant incremental improvement (permutation P = 0.096 and 0.074; Figure 6F). Cytotoxic–humoral covariation therefore identifies a biologically relevant association but not established predictive value beyond helper activity. Additional robustness and specificity analyses are provided in Figures S7E–K and S8E–I.

### An inferred T–B interactome nominates reciprocal antigen-presentation and helper pathways

To nominate molecular routes for investigating this participant-level coordination, we inferred candidate T–B communication pathways using MultiNicheNet [33,34] (Figure 6G). The selected pathways positioned B cells not only as recipients of T-cell help but also as potential antigen-presenting and costimulatory partners. B-to-T candidates included MHC-II–CD4 interactions involving Tfh and Th17 cells and CD86–CD28 interactions involving Treg or Th17 cells. Atypical-memory B cells featured prominently among selected antigen-presentation candidates, including HLA-DPB1–CD4 toward Tfh in inflamed UC and HLA-DRA–CD4 toward Th17 in the shared-series CD-versus-UC comparison (prioritization scores = 0.894 and 0.884). Reciprocal T-to-B candidates connected helper and regulatory populations to B-cell activation and plasma-cell states. These included Tfh/Th17 CD40LG–CD40, Treg TGFB1–TGFBR3 toward IgM-plasma cells in inflamed CD (score = 0.856), and Th17-derived BAFF (TNFSF13B) toward B-lineage receptors. In the displayed CD-versus-control context, the BAFF candidate linked Th17 cells to TNFRSF17 on switched-plasma cells. Together, these pathways suggest testable routes through which antigen-presenting B-cell states and helper/regulatory T-cell activity could participate in reciprocal immune communication in IBD.

### Repertoire sequence patterns carry diagnostic and clinical-state information

These biological analyses establish relationships between receptor architecture and cellular phenotype. We next assessed a complementary application of the resource: whether distributed repertoire sequence features contain information about diagnosis and contemporaneous clinical state. Participant-level models summarized short sequence patterns across TCR or BCR repertoires, testing distributed receptor features rather than individual disease-specific clonotypes (Figure 7A). Internally validated diagnosis models distinguished both CD and UC from controls, with median outer-fold TCR ROC AUCs of 0.989 and 1.000 and BCR AUCs of 0.844 and 0.821, respectively. Discrimination between CD and UC was lower, although still appreciable (TCR AUC = 0.889; BCR AUC = 0.739; Figure 7B). This pattern is compatible with a shared IBD-associated repertoire component alongside features distinguishing the two diseases, but classification alone cannot identify their biological origin.

**Figure 7.**
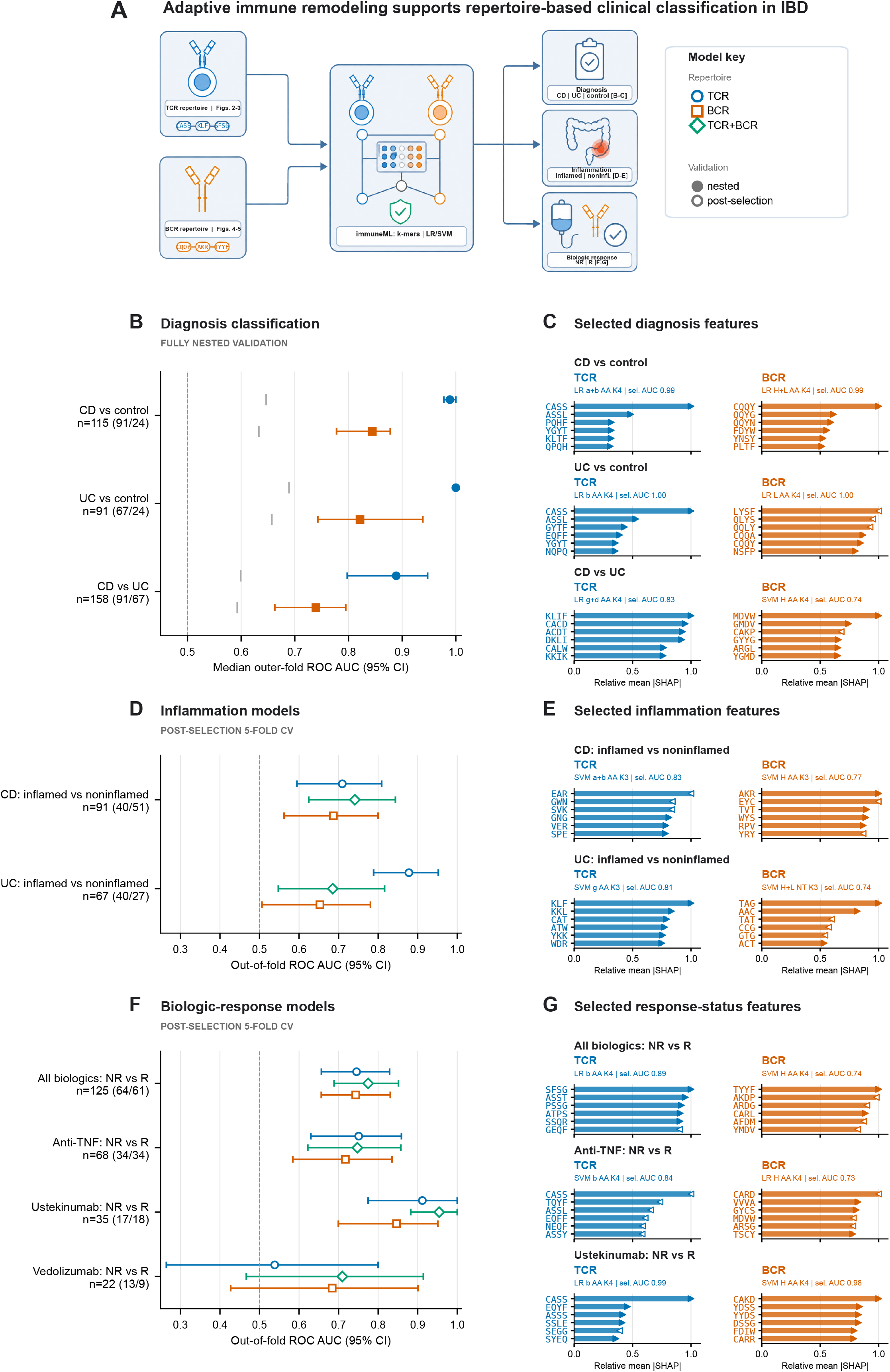
Repertoire benchmarks distinguish diagnosis and contemporaneous clinical state. (A) Repertoire-based clinical-classification workflow. (B) Nested participant-level diagnosis benchmarks using fixed abundance-weighted pooled α+β TCR or BCR heavy-chain amino-acid 4-mers. Points show median ROC AUC across 15 outer folds (five folds × three repeats); intervals bootstrap these correlated fold summaries. Participant totals and first-/second-class counts are shown beside each comparison in (B), (D), and (F). Gray ticks indicate the 95th percentile of fixed-pipeline permutation nulls. (C) Descriptive SHAP features from broader-screen-selected diagnosis models, which may differ from the primary models in (B). (D) Five-fold out-of-fold inflammation classification, conditional on prior endpoint-leader selection. Intervals show participant-bootstrap 95% CIs. (E) Selected inflammation-model SHAP features. (F) Conditional post-selection classification of contemporaneous six-month response status for pooled biologics, anti-TNF, ustekinumab, and vedolizumab. These are not pretreatment predictions. (G) Response-status SHAP features for pooled biologics, anti-TNF, and ustekinumab; vedolizumab attributions are provided in Source Data. In (C), (E), and (G), the six highest-ranked features are normalized within model. Filled rightward and open leftward markers indicate associations with the first and second labeled classes. Labels identify model, chain, sequence representation, k-mer length, and selection—not out-of-fold—AUC. Attributions derive from full-cohort refits and do not establish antigen specificity. The selected-model analyses in (C–G) use normalized k-mer frequencies from unique productive sequences, unlike the abundance-weighted primary benchmarks in (B). Combined-chain features pool sequences without preserving same-cell pairing. LR, logistic regression; SVM, support-vector machine; AA, amino acid; NT, nucleotide; H, heavy chain; L, light chain; a, b, g, and d denote α, β, γ, and δ chains. NR, nonresponder; R, responder. See also Figure S10 and Tables S5 and S6.

The primary diagnosis benchmarks used fixed sequence representations—pooled α+β TCR amino-acid 4-mers or BCR heavy-chain amino-acid 4-mers—with feature selection and model tuning confined to nested training folds. These estimates therefore assess internal participant-level generalization, not transfer to independently recruited populations. Calibration, threshold-dependent performance, permutation analyses, and selection optimism are documented in Figure S10; Tables S5 and S6 describe the broader model screen rather than the primary nested benchmarks.

Repertoire associations with active inflammation differed between diseases and receptor compartments. In CD, TCR, BCR, and joint models yielded ROC AUCs of 0.709, 0.687, and 0.741, respectively (n = 91). In UC, corresponding values were 0.878, 0.653, and 0.685 (n = 67; Figure 7D). Combining TCR and BCR features therefore did not consistently improve discrimination, despite the biological coordination observed between these compartments. Different sequence representations led the endpoint-specific screens, including α+β TCR features in CD and γ-chain features in UC. These differences nominate receptor compartments for further investigation but do not establish that a particular lineage drives intestinal inflammation. Moreover, pooled-chain representations do not preserve same-cell receptor pairing, and the clinical-state estimates were conditional on prior model selection within the same cohort. Associations with biologic-response status were similarly heterogeneous (Figure 7F). TCR, BCR, and joint AUCs were 0.745, 0.743, and 0.775 for pooled biologics (n = 125); 0.751, 0.717, and 0.747 for anti-TNF (n = 68); 0.912, 0.846, and 0.954 for ustekinumab (n = 35); and 0.538, 0.684, and 0.709 for vedolizumab (n = 22). Although the ustekinumab estimates were numerically high, the comparison included only 17 nonresponders and 18 responders, while vedolizumab estimates were particularly imprecise. Importantly, blood sampling coincided with the six-month response assessment.

Feature-attribution analyses identified short sequence patterns contributing to model-specific classification (Figure 7C,E,G). These patterns provide candidates for follow-up receptor mapping, but they are not equivalent to stable clonotypes or validated antigen-specific motifs. Attribution models were fitted to the full cohort, and their displayed selection AUCs are distinct from out-of-fold performance; diagnosis attribution models also need not match the fixed primary benchmark models. Together, these analyses establish a framework for evaluating circulating repertoire features as candidate markers of diagnosis and clinical state. Their translational significance lies in the possibility of reading systemic immune remodeling from blood. Prospective, independently sampled cohorts will be required to distinguish persistent immune history from current inflammatory burden and treatment-associated changes, and to determine whether repertoire profiling adds useful information beyond established clinical measures.

## DISCUSSION

This resource connects circulating adaptive immune-cell states to complementary features of the receptors expressed by those cells. Clonal expansion was associated with inflammatory specialization within matched T-cell states. Paired-receptor relatedness linked similar programs across participants, partly through shared cellular identity. Accumulated BCR mutation, measured by divergence from germline, was associated with selected plasma-cell transcriptional programs, whereas sequence diversification among sampled members of the same B-cell lineage was not significantly increased in IBD. T-cell programs that support B-cell responses remained associated with specific B-cell states after accounting for other measured factors, consistent with coordinated T- and B-cell responses. Together, these findings show that clonal expansion, receptor sequence similarity, and accumulated mutation provide distinct information about immune-cell states. These relationships would be obscured by describing the adaptive immune response simply as more diverse, more expanded, or more activated.

A central T-cell finding is that larger clones were not simply more abundant representatives of otherwise equivalent cells. Within participants and annotated states, increasing clone size was associated with stronger GZMK inflammatory-memory, EOMES–ZEB2, cytotoxic, and Th1/Tc1 programs. Larger conventional-CD8 clones also occupied later positions along the transcriptional continuum while spanning a broader range of states. Their expansion was therefore compatible with both effector specialization and the maintenance of transcriptional diversity within a clone. This may be important in chronic inflammation, where cells sharing a receptor can occupy different functional states rather than converge on a single terminal phenotype. The external UC analysis sharpened the biological interpretation of this relationship. Clone size correlated positively with cytotoxic, GZMK inflammatory-memory, and pathogenic-Th17 programs, but negatively with conventional-Th17 and naive/central-memory programs [28]. This argues for separating inflammatory effector features from conventional Th17 identity when interpreting expanded circulating T cells in IBD. It does not imply that expansion excludes the Th17 lineage: the pathogenic-Th17 signature includes genes shared with cytotoxic and inflammatory programs, and the external analysis was not state matched. The internal focused Th17/Treg analyses likewise did not establish significant expansion-associated increases across the tested programs. Together, these observations favor a description based on functional programs over a broad claim of “Th17 expansion,” while preserving the possibility that distinct Th17 populations have different roles in disease.

Receptor similarity provided a second view of this organization. Non-identical paired αβ TCRs carried concordant inflammatory and cytotoxic programs across participants, including among singleton clonotypes. The relationship was therefore not restricted to large clones or identical receptors shared between individuals. Its attenuation after cell-state adjustment is equally informative: sequence-related receptors were partly associated with similar cellular identities, rather than exerting a demonstrably independent influence on transcription. The additional organization of neighborhoods by current inflammatory status connects this receptor–state relationship to clinical context. Shared antigen exposure is one possible explanation, but receptor-generation biases, selection history, and common inflammatory environments may also contribute. Previous sequence-clustering studies and evidence for microbiota- and yeast-reactive T cells in IBD provide a basis for experimental follow-up [12,13,21,25,29,30]. The paired-chain neighborhoods identify specific receptors for such testing without assigning a common antigen.

The B-cell findings point to accumulated maturation rather than a uniform increase in antibody-producing activity. Heavy-chain mutation was elevated in both diseases, with preferential CDR targeting and the clearest isotype-matched increase in IgM. The involvement of IgM indicates that the maturation-associated signal is not confined to class-switched receptors. Lineage reconstruction further separated distance from germline from diversification among the sequences present at sampling. Greater germline-inclusive branch length without significantly greater observed-only divergence is consistent with the circulation containing more extensively mutated receptors but does not establish an increased current mutation rate. In UC, the association between CDR targeting and fecal calprotectin provides an additional link between systemic receptor maturation and intestinal inflammatory burden, although this relationship was not detected in CD.

Transcriptional analyses helped distinguish this maturation signal from clonal expansion. Expanded B cells did not show a common increase in every secretory program. Antibody-secretion, IgA-associated, and plasma-differentiation features differed from IgG-inflammatory and antigen-presentation-associated programs. At the lineage level, IgA-mucosal and plasma-differentiation activity remained associated with mutation burden and class-switching probability after accounting for participant and annotated state and excluding constant-region genes. Neither program was associated with every measure of lineage complexity. These selective relationships suggest that aspects of plasma-cell specialization accompany accumulated receptor maturation without requiring broader observed diversification or occupancy of multiple states. They also explain why lineage-level associations can coexist with nonsignificant adjusted diagnosis-level switching differences. In the context of microbial antibody recognition and disease-associated B-cell clones [24,26,27], the atlas offers a way to connect candidate antibody functions to the cellular programs and maturation histories of the cells carrying them.

The cross-compartment findings place these receptor-associated states within a coordinated immune response. Tph/Tfh-help activity retained independent associations with IgG-inflammatory plasma, atypical-memory, and antigen-presentation programs, while suppressive-Treg activity contributed independently to B-cell antigen presentation. Regulatory and inflammatory programs can therefore covary across individuals. Their coexistence should not be taken as evidence that regulation is absent or effective. It may reflect compensatory activity, shared stimulation, or parallel responses to disease. The relative scarcity of mixed Th17/Treg clonotypes further shows that coordination across populations need not depend on abundant shared clones. In CD, expanded-TCR cytotoxicity also tracked IgA-mucosal and plasma-differentiation activity, although it did not significantly improve held-out performance beyond helper activity. This association may reflect a common inflammatory setting rather than direct cytotoxic T-cell control of plasma differentiation.

The inferred interactome suggests molecular routes through which some of this coordination could occur [33,34]. B-cell MHC-II and CD86 candidates position antigen-presenting B-cell states alongside Tfh, Th17, and Treg populations, while reciprocal CD40L, BAFF, and TGF-β candidates connect T-cell activity to B-lineage states. The prominence of atypical-memory B cells among selected antigen-presentation candidates is notable because their potential role would not be captured by antibody output alone. Likewise, the appearance of IgM- and switched-plasma states among candidate receivers raises questions about how helper and regulatory inputs accompany humoral specialization. These are hypotheses derived from expression and prior interaction knowledge, not independent confirmation of the program correlations. The selected pairs provide focused candidates for perturbation and receptor-specific studies.

Peripheral blood offers access to this systemic organization, but it is not a direct substitute for intestinal tissue. Circulating cells may retain features acquired during tissue exposure, respond to systemic inflammation, or reflect treatment-associated changes. Our external blood–mucosal comparisons did not resolve these alternatives or establish trafficking direction [17,32]. A complementary longitudinal perspective [35] reported changes in microbiota-directed antibody binding, repertoire–microbiome relationships, and tolerogenic or gut-homing programs during a vitamin D intervention in IBD. Those findings motivate asking whether the receptor-associated states identified here are modifiable and whether their changes accompany altered microbial recognition. They do not independently validate the present clonotypes or inferred interactions. Longitudinal blood–tissue sampling will be needed to determine which circulating features track ongoing mucosal activity and which persist as a record of prior immune experience.

Clinical classification provides a complementary demonstration of the information contained in circulating repertoires. Strong internal diagnostic discrimination suggests that distributed sequence patterns could support repertoire-informed patient stratification, while associations with inflammation and biologic-response status motivate longitudinal studies of treatment-associated immune remodeling. Independent validation must establish whether repertoire profiling adds information beyond established clinical and inflammatory measures. Planned release of participant-level source data, receptor definitions, frozen model inputs, and fold assignments, together with the publicly available analysis code, will enable reproducible external testing and support development of these biological associations into clinically evaluable tools. By linking receptor architecture to cellular phenotype, this resource enables the selection of specific receptors and immune states for subsequent investigation. Paired-TCR neighborhoods nominate receptors for testing shared recognition. BCR lineage and program annotations guide antibody studies that distinguish accumulated maturation from functional specificity, and helper–B-cell associations prioritize candidate cellular relationships for perturbation. Longitudinal and paired blood–tissue studies can then determine which circulating features track mucosal activity and which persist after inflammation resolves. These applications build on the observed organization of adaptive immunity without assuming that receptor similarity establishes antigen specificity or that coordinated programs demonstrate direct cellular interaction.

### Limitations of the study

This resource is observational, primarily cross-sectional, and restricted to peripheral blood. Treatment exposure, age, disease location, inflammatory burden, and sequencing depth may confound repertoire structure, although our sensitivity analyses minimize these considerations. Correlations across participants do not establish direct cellular interaction or mediation. Expansion does not imply pathogenicity and sequence neighborhoods do not establish antigen specificity. In addition, pseudotime does not establish temporal differentiation, and BCR lineages do not prove the direction of class switching. The machine-learning results are research benchmarks rather than prospective diagnostic, monitoring, or treatment-response tests. Independent IBD and gastrointestinal disease-control cohorts, prespecified pipelines, longitudinal sampling before and after therapy, and paired blood-intestinal measurements are required. Recombinant receptor expression, peptide-MHC screening, microbial antigen libraries, phage display, and antibody-binding assays will be needed to establish specificity and function. These studies can determine whether the circulating states identified here precede relapse, change with mucosal healing, or define mechanistically distinct endotypes.

## TABLE

Unless stated otherwise, CD denotes Crohn’s disease, UC denotes ulcerative colitis, and control denotes non-IBD controls. Participants are the biological replication unit; cell, clonotype, lineage, and eligible-participant counts are distinguished in the legends and STAR Methods. FDR denotes false discovery rate; CI, confidence interval; IQR, interquartile range. Clinical definitions are provided in STAR Methods and cohort characteristics in Table S1.

## Supplemental information

### Supplemental figure legends

**Figure S1.**
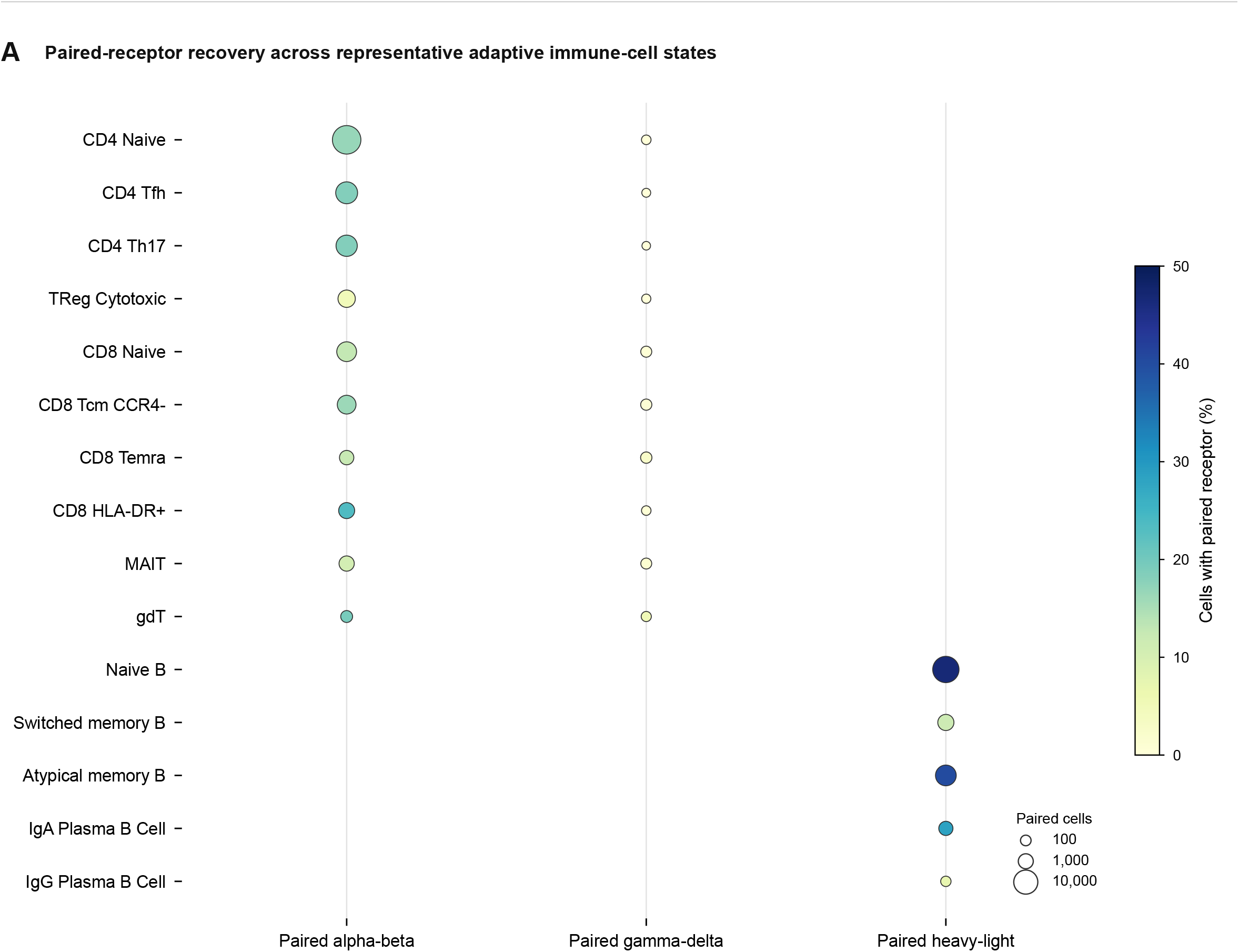
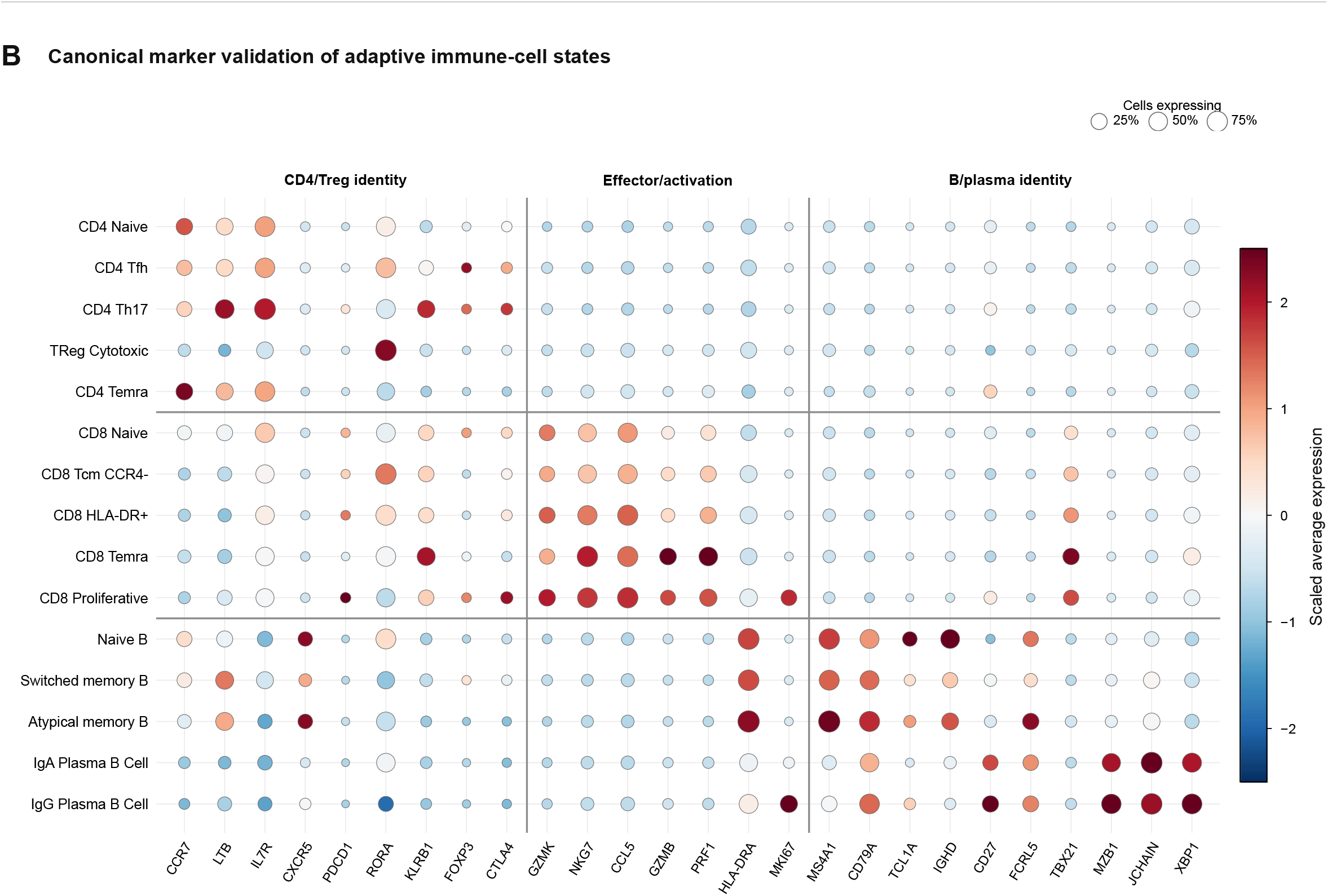
Receptor recovery and adaptive-state validation. (A) Productive paired alpha-beta, paired gamma-delta, and paired heavy-light receptor recovery across representative adaptive immune-cell states; dot size reports paired-cell number and color reports the percentage of cells within each state. (B) Canonical marker validation across representative CD4/Treg, CD8 T-cell, B-cell, and plasma-cell states; dot size indicates the percentage of cells expressing each gene and color indicates gene-wise scaled average expression. Both panels use the complete quality-controlled lineage datasets.

**Figure S2.**
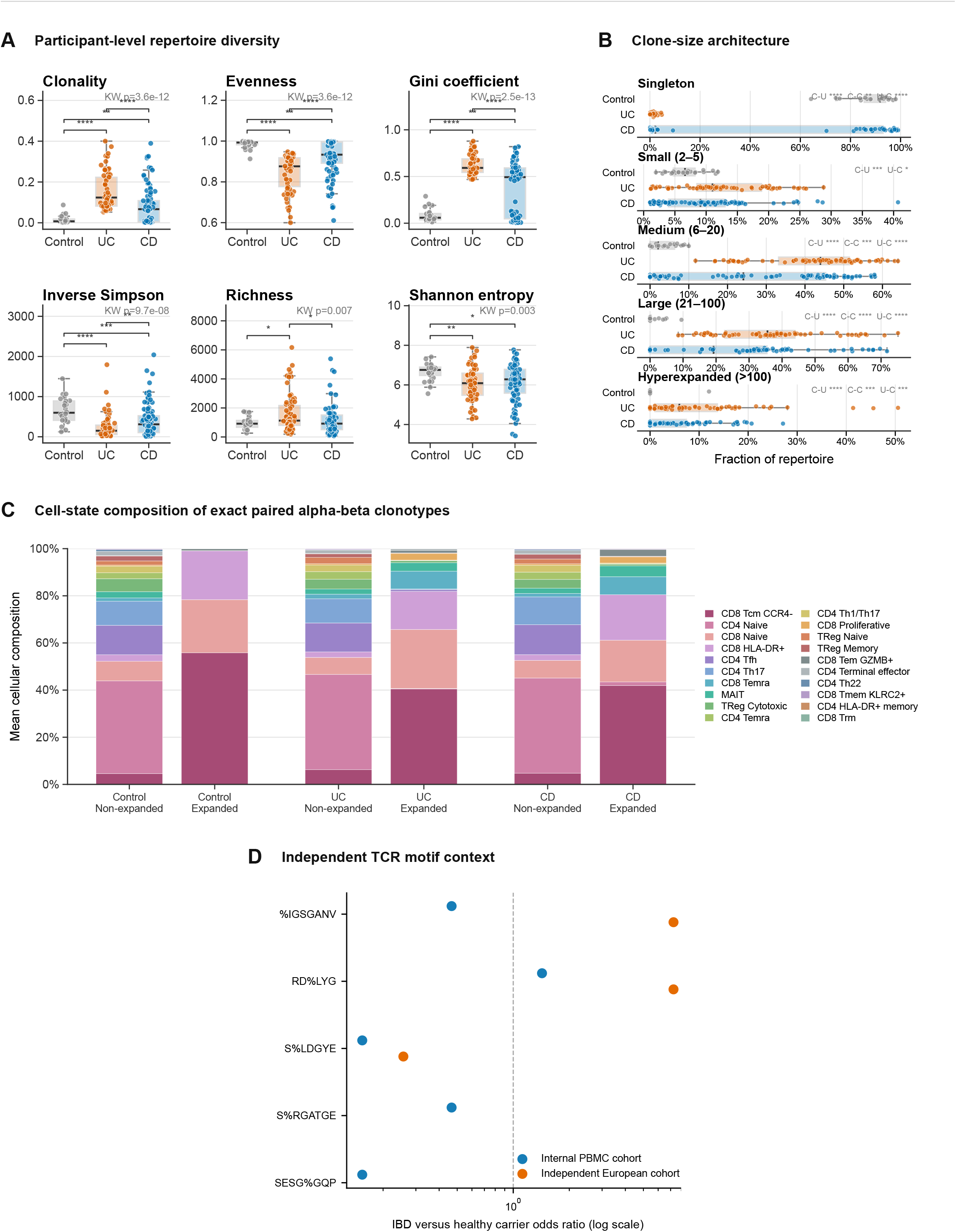
TCR repertoire structure and external motif context supporting Figures 2 and 3. (A) Participant-level diversity and clonality. (B) Diagnosis-stratified clone-size composition. (C) Cell-state composition of non-expanded and expanded exact paired alpha-beta clonotypes. (D) Carrier odds ratios for five previously reported GLIPH2 specificity motifs in the internal PBMC cohort and an independent European cohort. The external motif comparison supports the general existence of convergent IBD-associated motifs but does not validate the manuscript-specific sequence groups.

**Figure S3.**
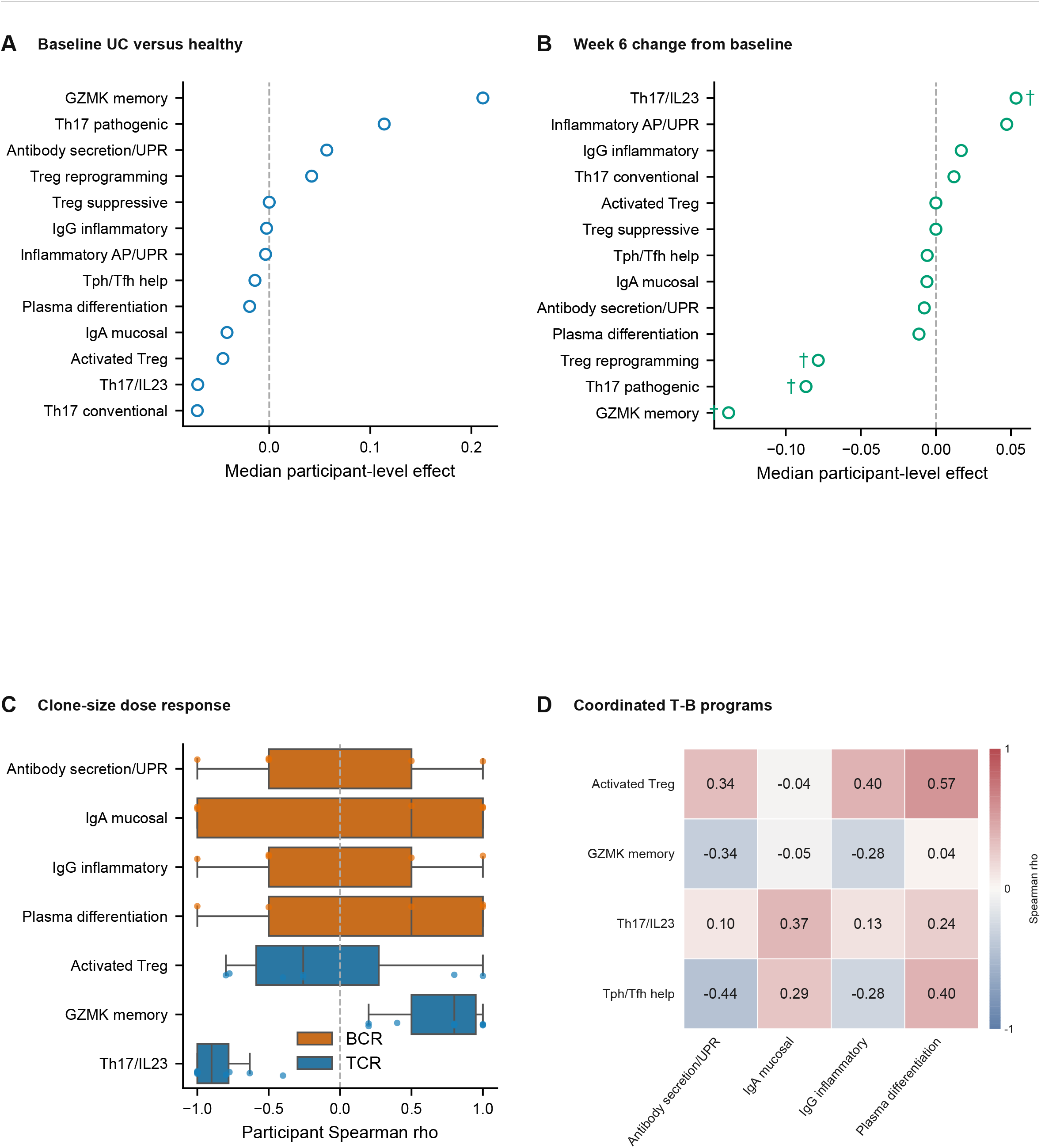
Independent longitudinal PBMC assessment of clone-associated programs, related to Figure 2. Public GSE261334 data [28] included five healthy participants and ten participants with UC sampled at baseline and week 6. (A) Baseline UC-minus-healthy program-score differences. (B) Paired week-6-minus-baseline changes in UC. Neither family contained FDR-significant effects. (C) Participant-level Spearman correlations between ordered exact paired-clone-size bins and program score for selected T- and B-cell programs; complementary TCR programs are shown in Figure 2H. (D) Baseline participant-level T–B program correlations (n = 15); no association passed FDR correction. Points in (A,B) show median effects without confidence intervals; daggers denote 0.05 ≤ FDR <0.10. Boxes in (C) show participant medians and IQRs. Positive or negative clone-bin slopes are associations, not evidence that expansion selects or excludes entire cell lineages. Longitudinal changes were not tested as response effects because participant-linked response labels were unavailable for reanalysis.

**Figure S4.**
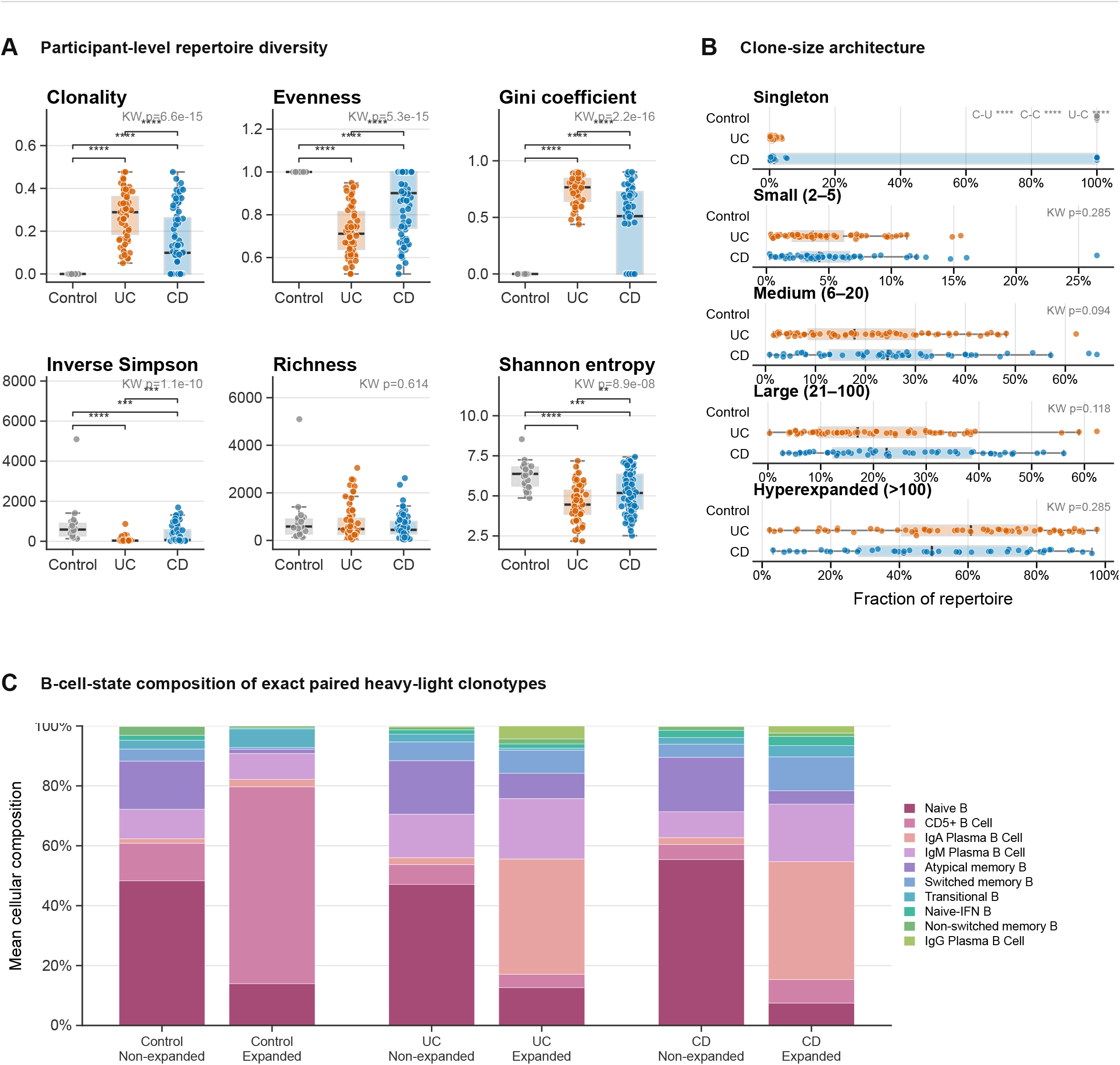
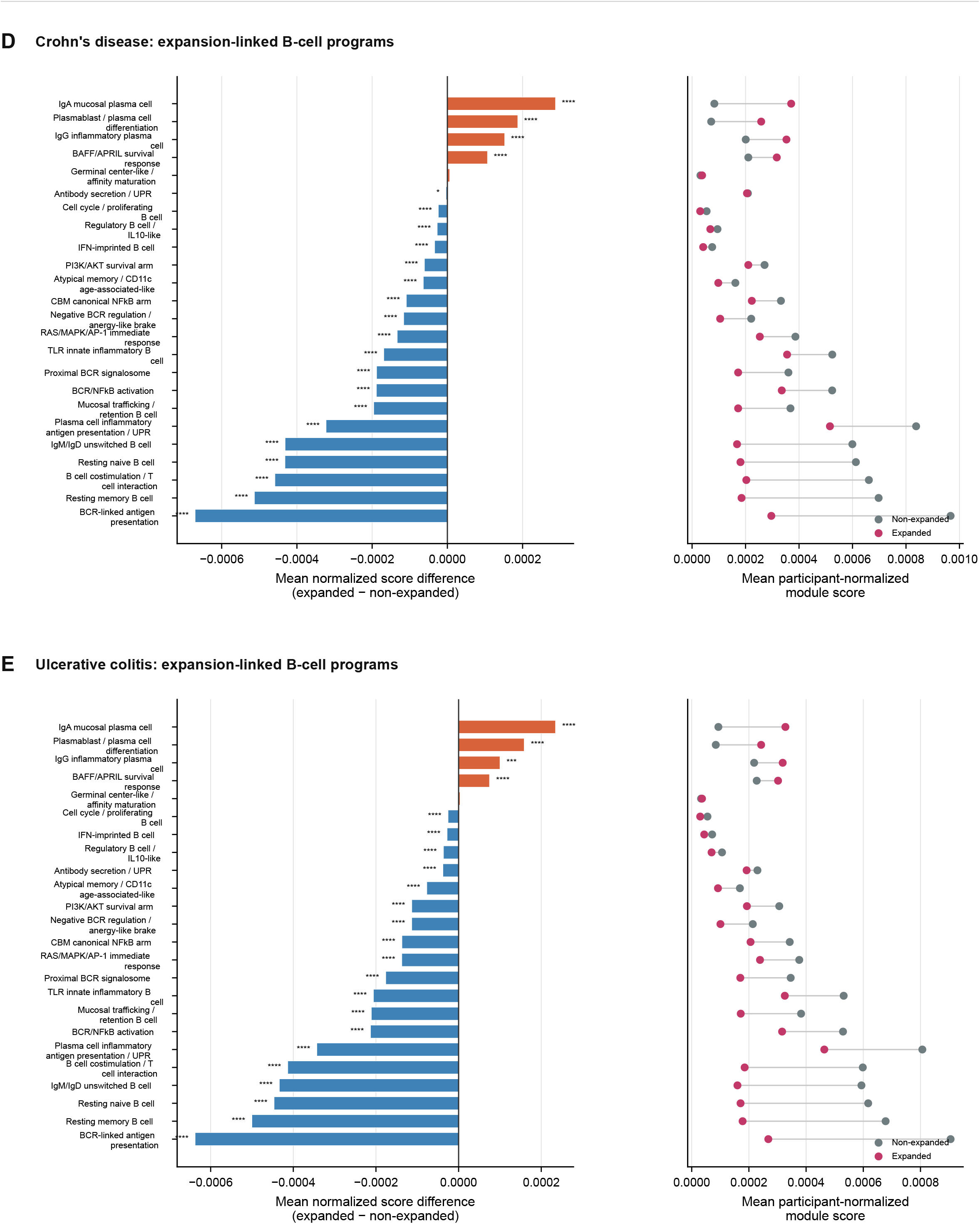
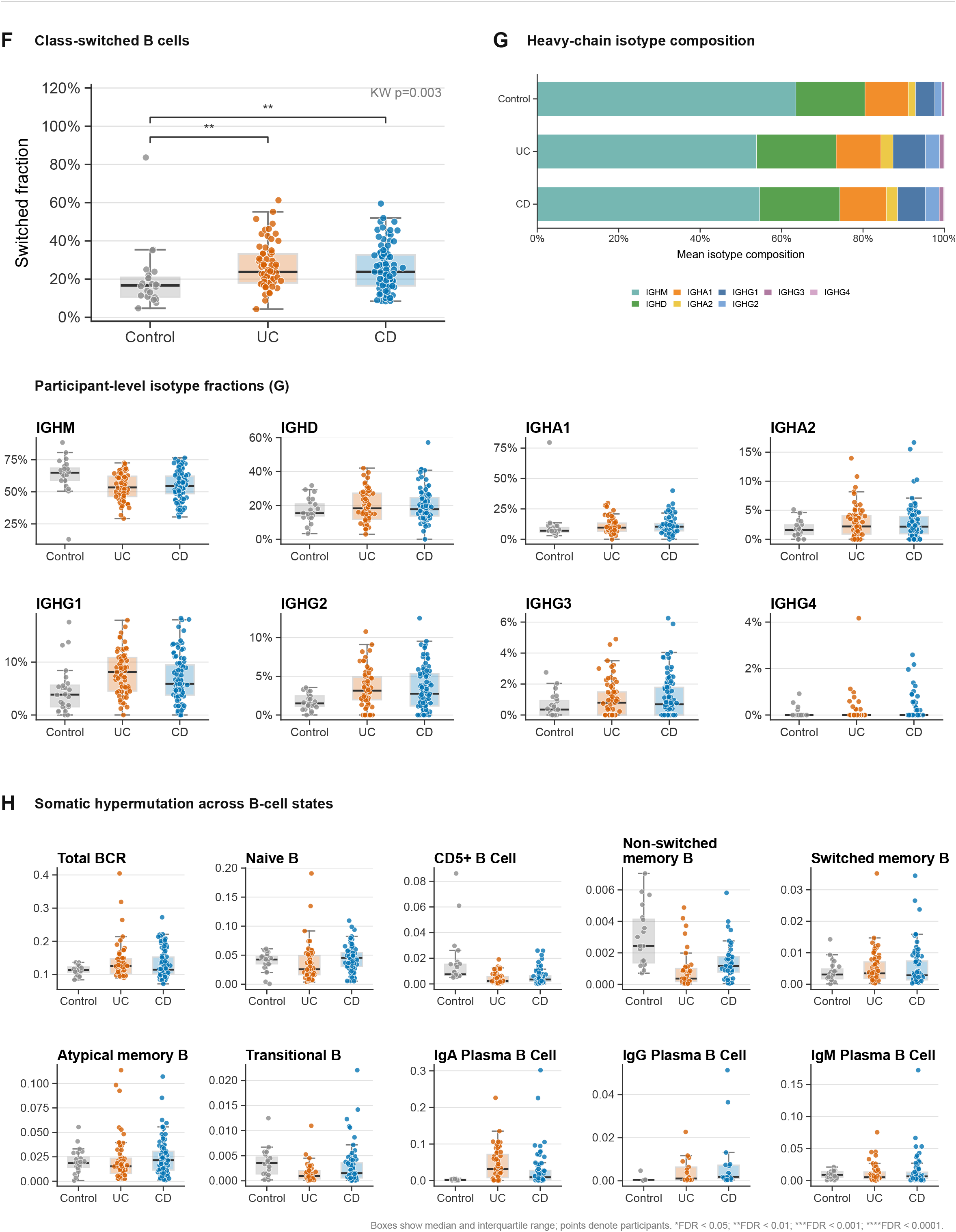
BCR repertoire structure, expansion-associated programs, isotype, and SHM, related to Figures 4 and 5. (A) Participant-level diversity and clonality calculated directly from positive-count records in the IBDBCRonly source dataset. These row-based summaries differ from the CDR3-amino-acid-aggregated immunarch clonality shown in Figure 4A; their numerical values are not interchangeable. (B) Diagnosis-stratified clone-size composition. (C) State composition of expanded and nonexpanded paired-chain BCR clonotypes. (D,E) CD- and UC-stratified legacy expanded-versus-nonexpanded program summaries, respectively; these are complementary to, not replacements for, the heavy-chain-defined state-matched pseudobulk contrasts in Figure 4C,D. (F) Class-switched fractions. (G) Diagnosis-level mean isotype composition and participant-level isotype fractions. (H) Heavy-chain SHM within matched B-cell states. Observed isotype fractions should be interpreted alongside the adjusted count models in Figure 4G. These analyses do not establish antigen specificity or an expansion-driven increase in plasma programming.

**Figure S5.**
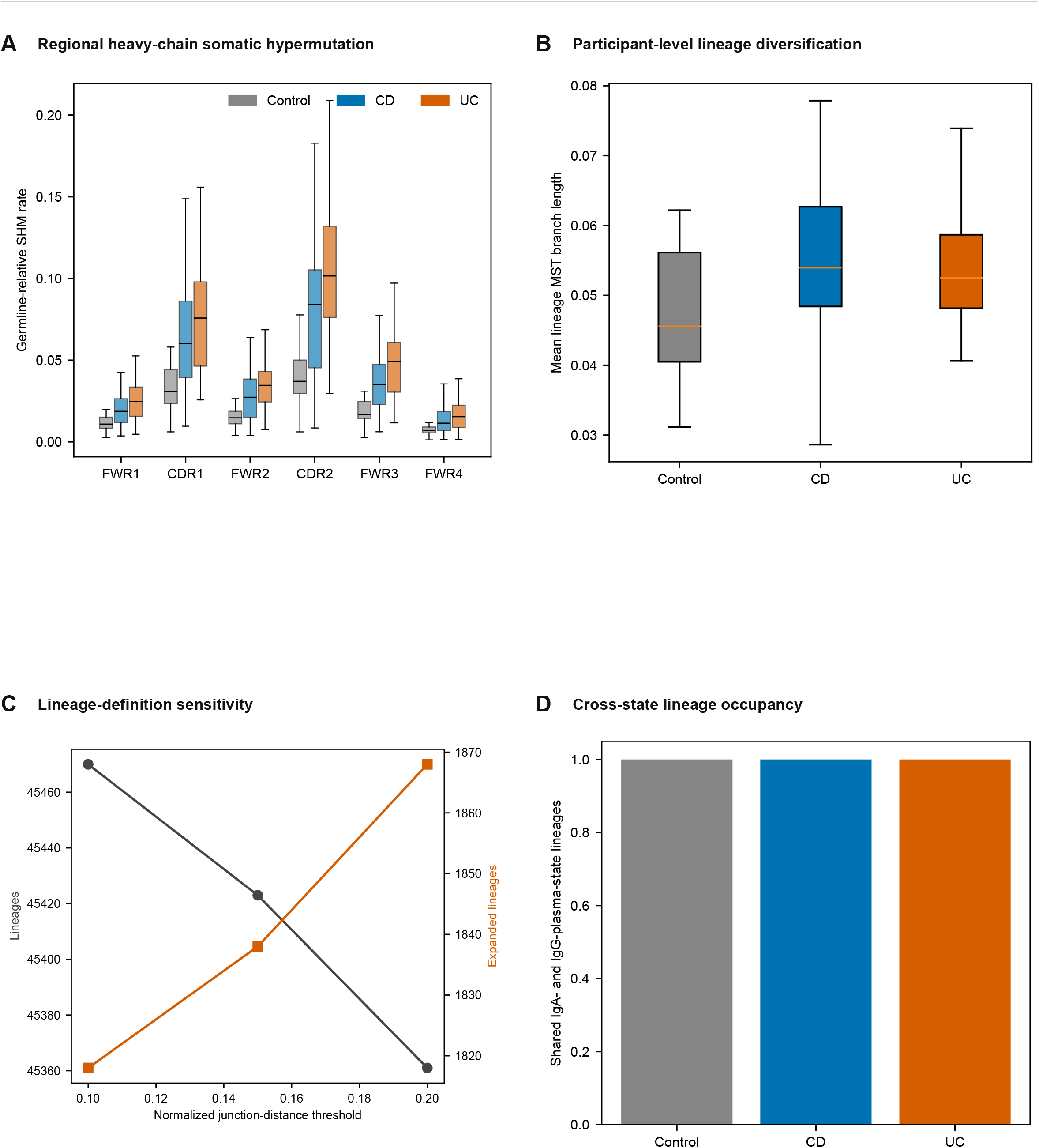
BCR germline-lineage reconstruction and definition sensitivity, related to Figure 5. (A) Participant-level regional heavy-chain SHM across framework and complementarity-determining regions. (B) Participant-level mean lineage minimum-spanning-tree branch length. (C) Sensitivity to normalized junction-distance thresholds of 0.10, 0.15, and 0.20. (D) Counts of heavy-chain lineages linked to both IgA-plasma and IgG-plasma transcriptional states: one detected lineage per diagnosis. These are detected lineage counts, not fractions, observed class-switch events, or estimates of trafficking. Light-chain-linked subdivisions and limited cell-to-full-heavy-sequence resolution constrain interpretation. The analyses distinguish accumulated germline distance from diversification among observed descendants.

**Figure S6.**
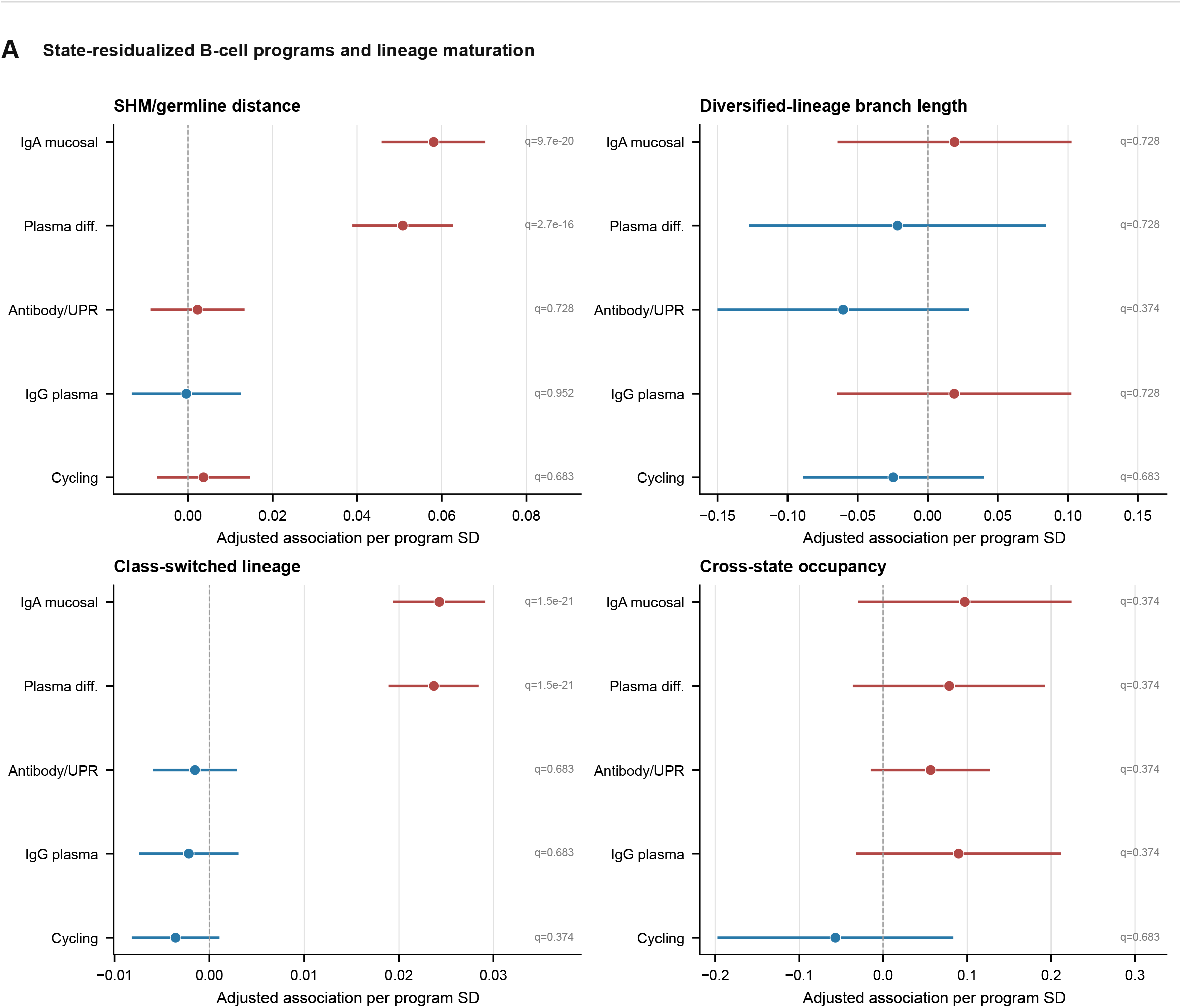
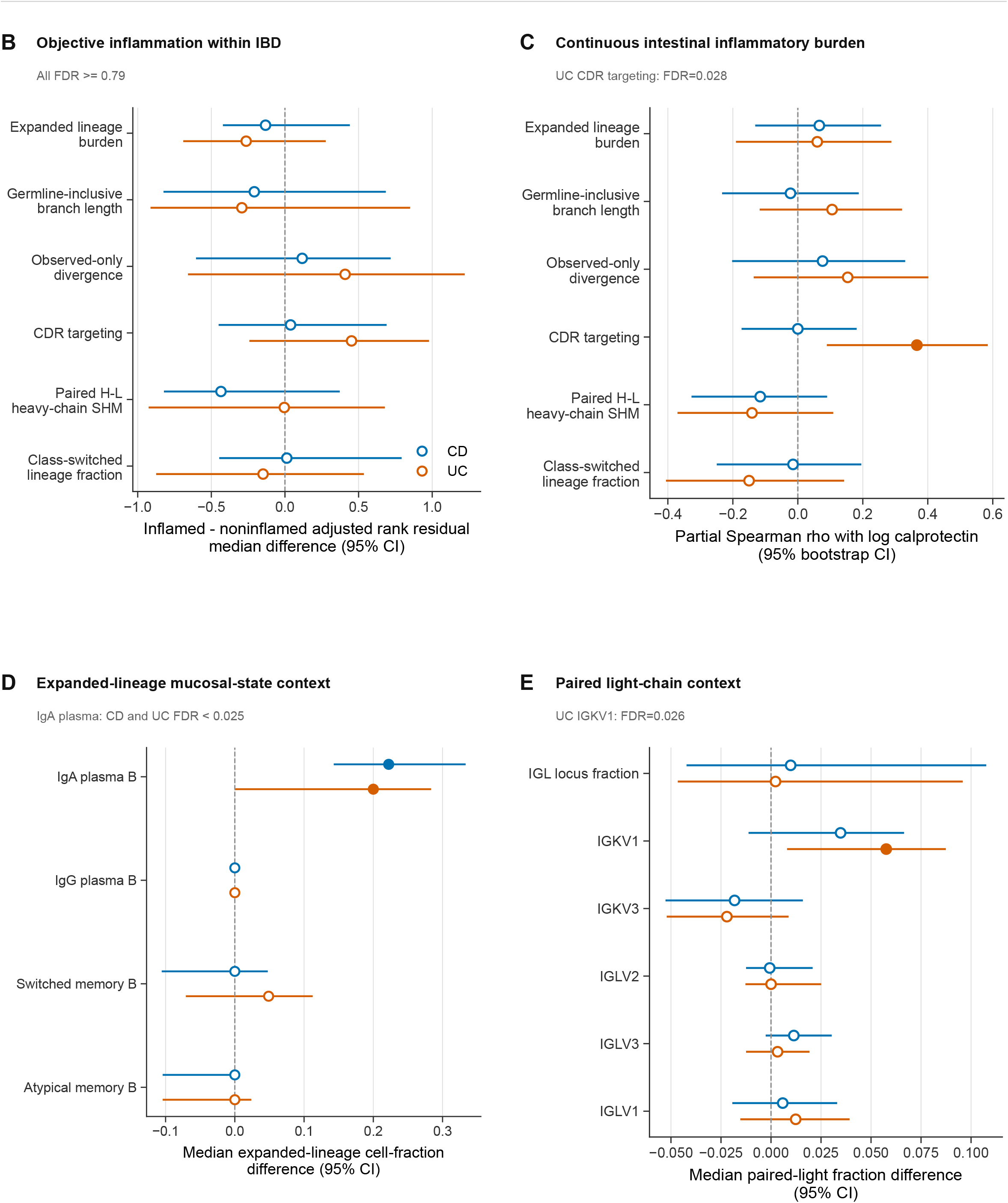
BCR lineage maturation and clinical context, related to Figure 5. (A) Participant-aware associations of five state-residualized, constant-region-excluded programs with germline distance, branch length among diversified lineages, class-switching probability, and cross-state occupancy. Continuous outcomes are standardized; switching effects are absolute probabilities. Models adjusted for lineage abundance, scored-cell depth, and pseudotime, with participant fixed effects and participant-clustered errors; FDR was controlled across 20 tests. (B) Objective-inflammation contrasts. (C) Associations with continuous fecal calprotectin. (D) Mucosal-state composition of expanded lineages. (E) Paired-light-chain context. Points and intervals denote the corresponding participant-aware estimates and 95% CIs where shown. In (B–E), blue and orange denote CD and UC, respectively; filled symbols indicate FDR <0.05. These cross-sectional analyses do not establish mutation rate, temporal switch direction, antigen specificity, or an effect of exact-clonotype expansion on plasma programming.

**Figure S7.**
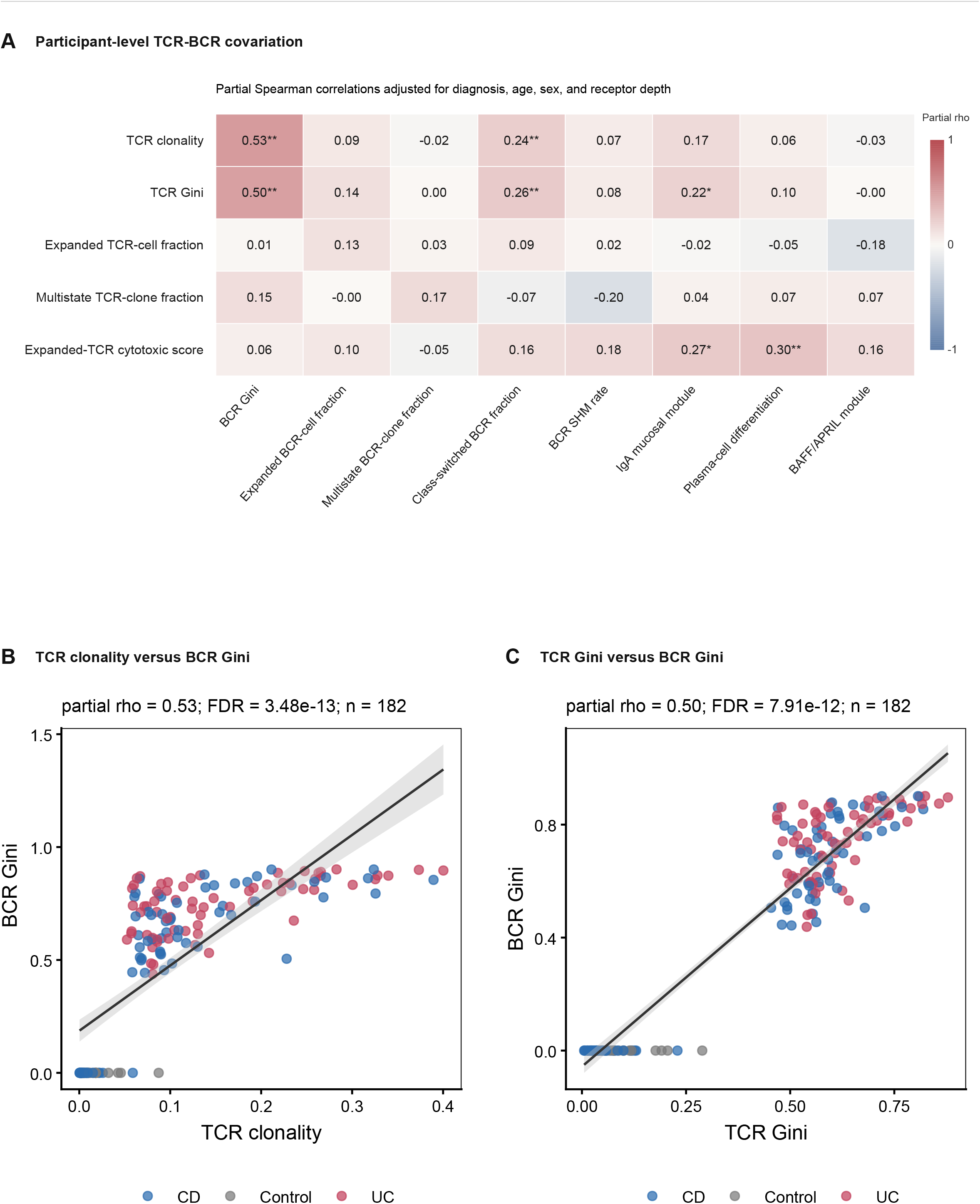
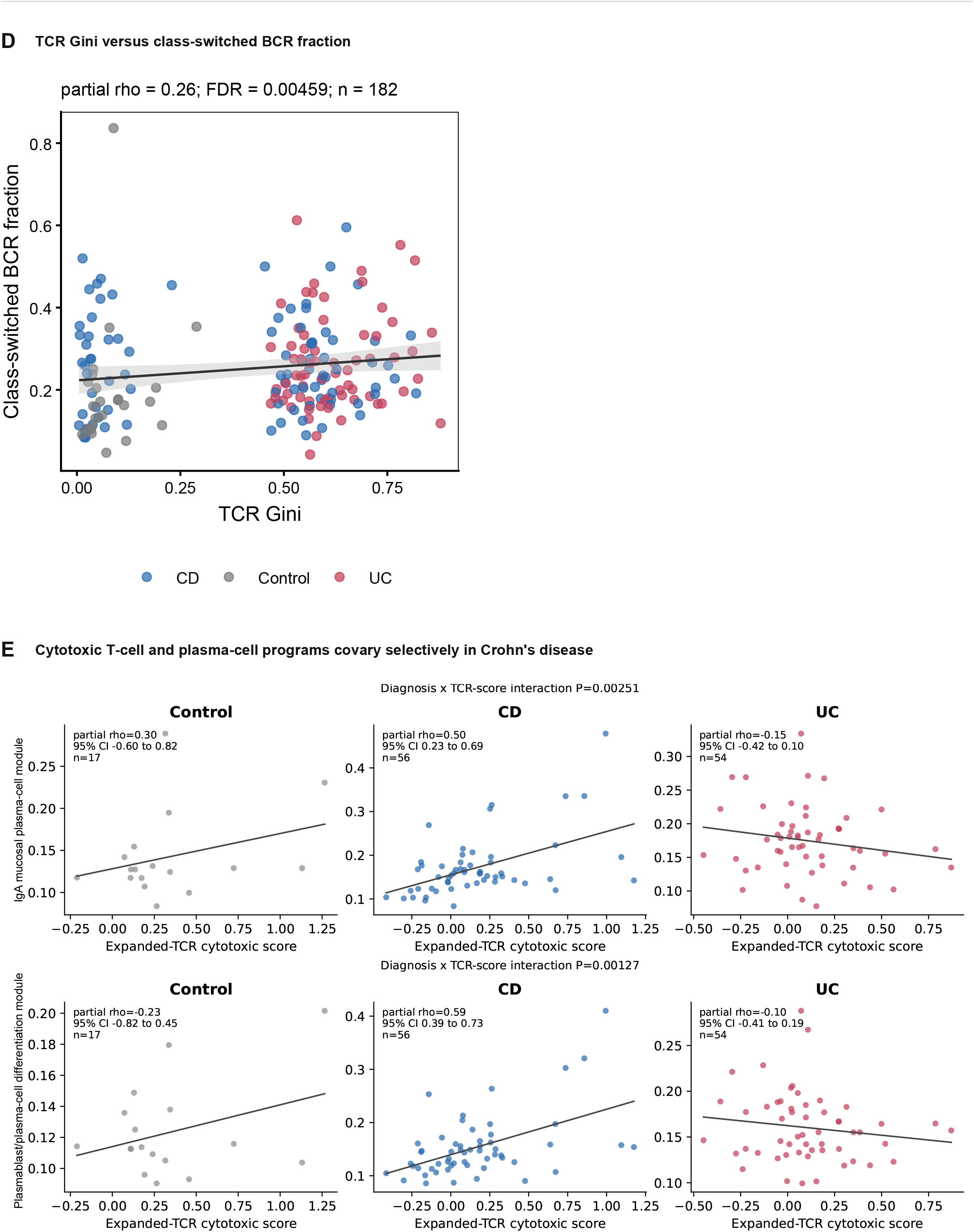
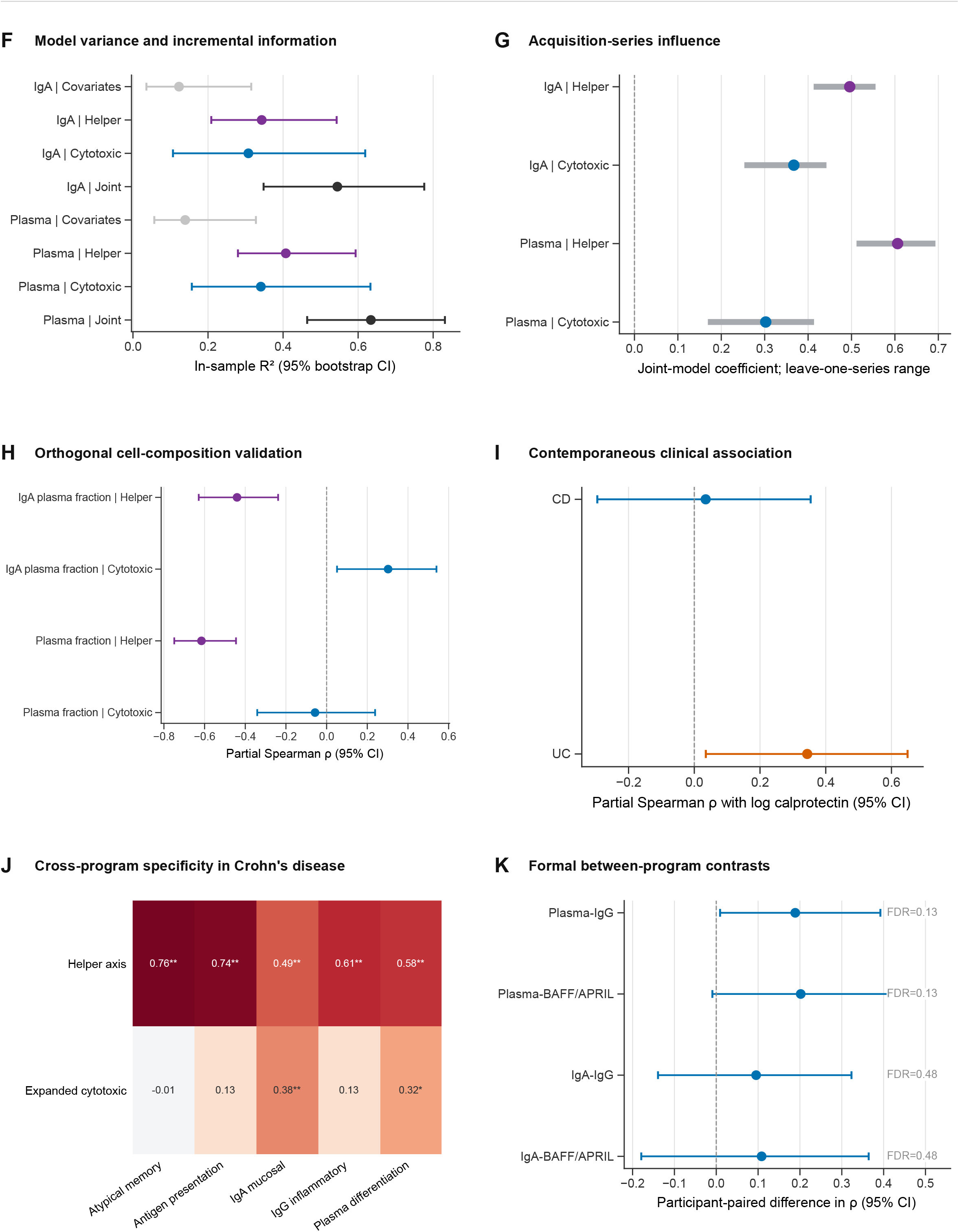
Global TCR–BCR coordination and robustness, related to Figure 6. (A) Complete participant-level cross-compartment partial-correlation screen. (B–D) Representative repertoire associations; scatterplots show unadjusted participant values, whereas annotations report covariate-adjusted partial correlations. (E) Diagnosis-stratified cytotoxic–plasma program associations. (F) In-sample model variance and incremental information; these R² summaries differ from the held-out correlations in Figure 6F. (G) Leave-one-acquisition-series influence. (H) Cell-composition sensitivity. (I) Contemporaneous clinical associations. (J) Cross-program specificity in CD. (K) Formal between-program contrasts, none significant. In (A,J), asterisks indicate FDR <0.05 (*) or <0.01 (**). Covariates and resampling are specified in STAR Methods. The analyses assess covariation, not direct interaction or mediation.

**Figure S8.**
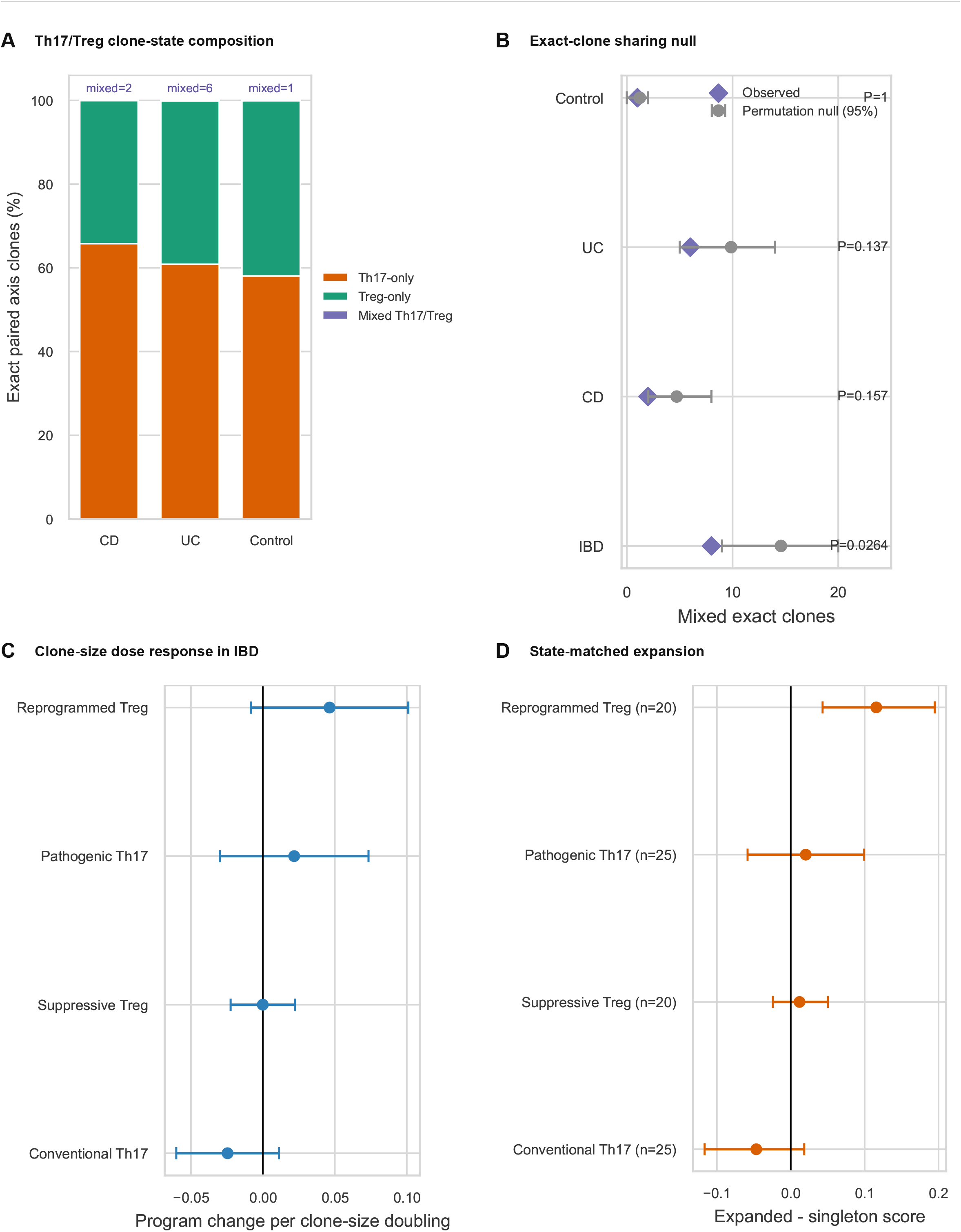
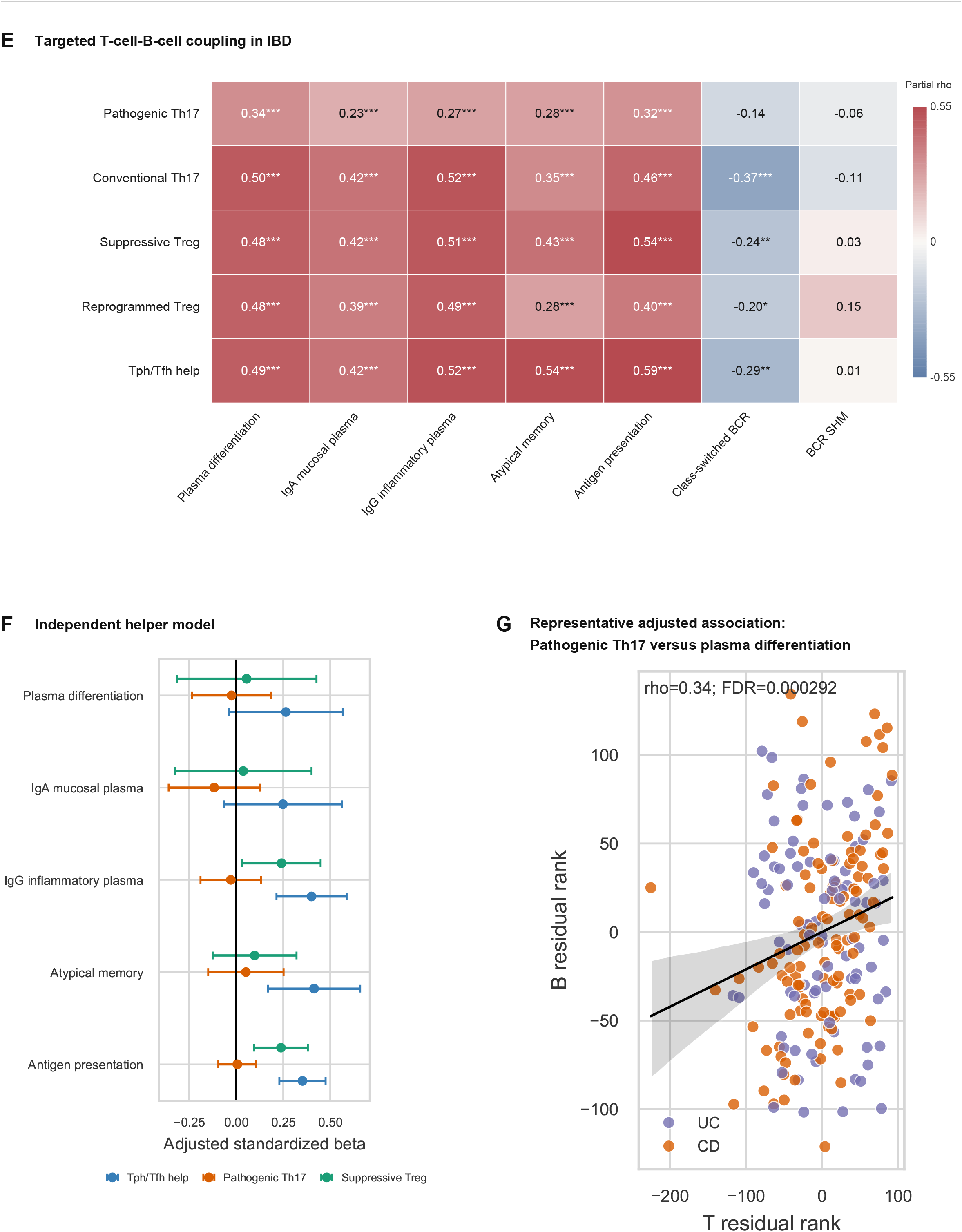
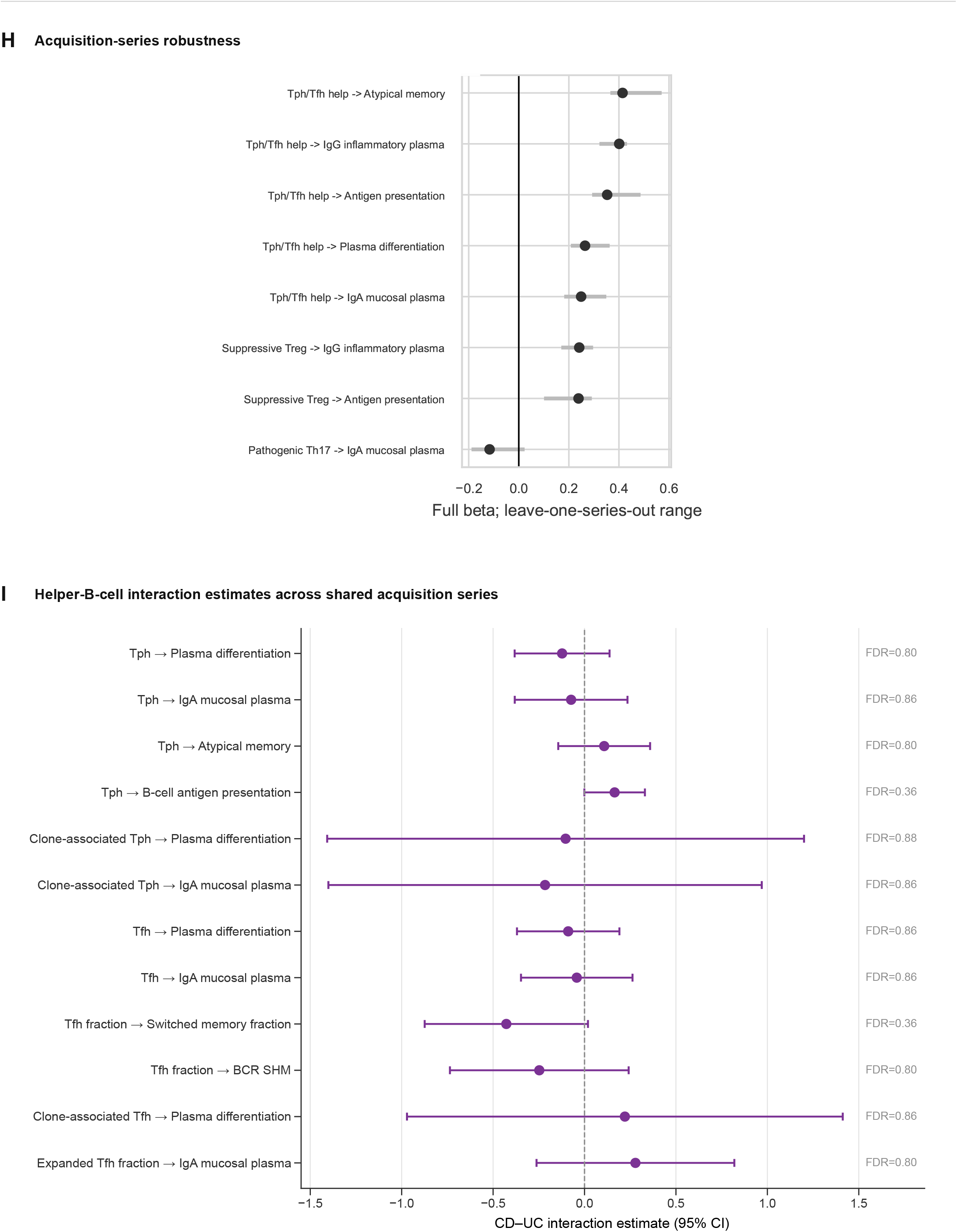
Helper, Th17, and Treg clone architecture and B-cell coordination, related to Figure 6. (A) Exact paired αβ Th17/Treg clone composition. (B) Within-participant state-label null for mixed clones (10,000 permutations); eight observed mixed IBD clones versus 14.57 expected on average. Gray intervals show the central 95% of permutation-null counts; diamonds mark observed counts. (C) Clone-size effects. (D) State-matched expanded-minus-singleton program contrasts. (E) Participant-level helper/regulatory–B-cell partial correlations; asterisks indicate FDR <0.05 (*), <0.01 (**), or <0.001 (***). (F) Independent standardized joint-model contributions. (G) Representative adjusted pathogenic-Th17–plasma-differentiation association. (H) Leave-one-acquisition-series sensitivity. (I) CD-versus-UC interactions restricted to shared series; none passed FDR correction. Clone-size and expansion contrasts for the focused Th17/Treg programs were not FDR significant. Mixed-clone depletion distinguishes clone sharing from participant-level coordination but does not establish the mechanism of that coordination.

**Figure S9.**
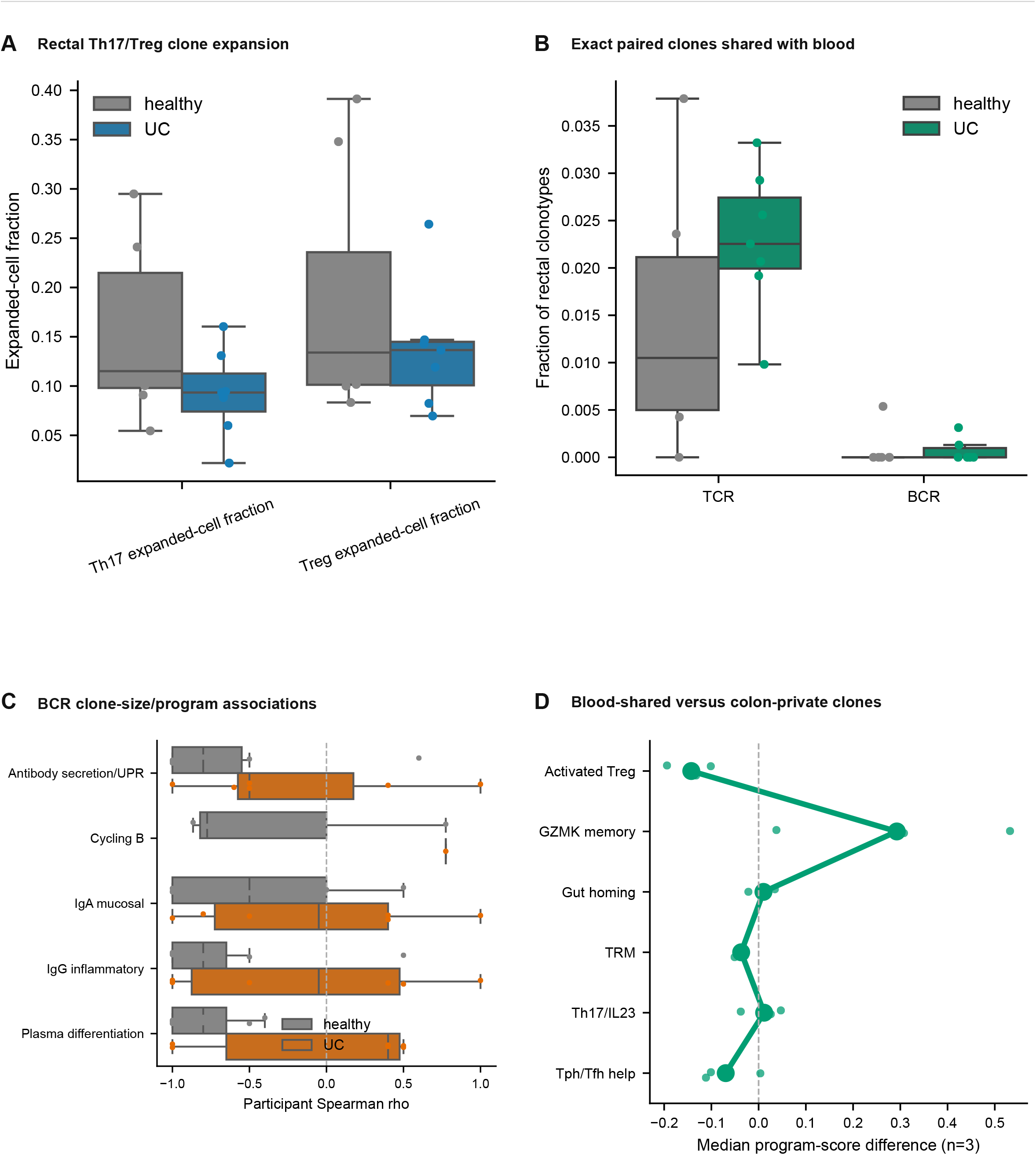
External mucosal and blood–tissue analyses, related to Figures 2, 5, and 6. (A) Rectal Th17-like and Treg expanded-cell fractions in GSE125527 [17]. (B) Exact paired TCR or BCR clonotypes shared with blood. (C) BCR clone-bin/program associations in the same external resource. Disease comparisons did not pass FDR correction. (D) Program-score differences between blood-shared and colon-private TCR clonotypes in three CD participants from GSE301689 [32]. These n = 3 estimates are descriptive, and none passed FDR correction. Cross-tissue identity does not establish trafficking direction; the public BCR files lacked the germline-alignment information needed to validate the SHM findings.

**Figure S10.**
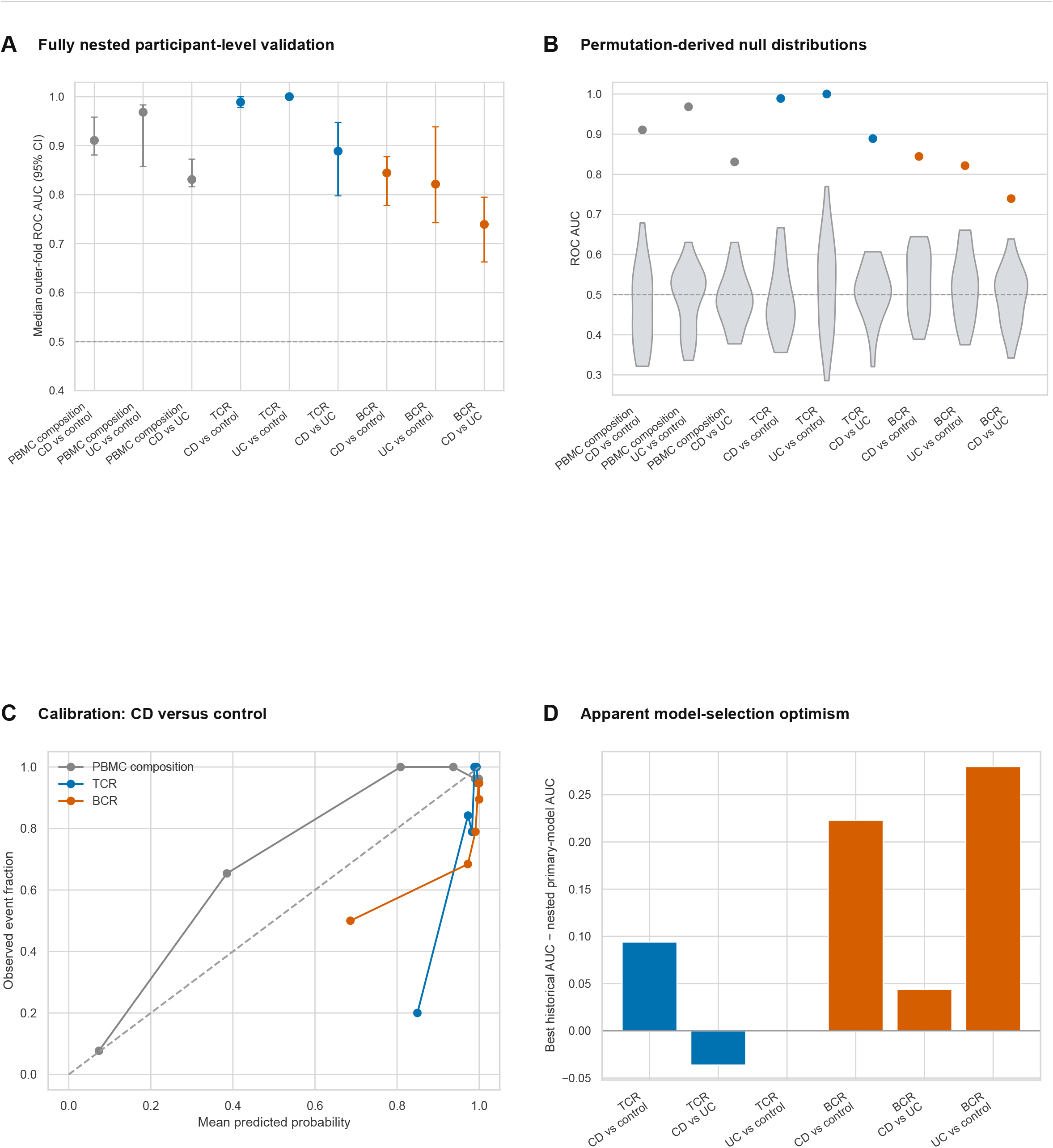
Diagnosis benchmarking, calibration, and selection optimism, related to Figure 7. (A) Nested participant-level performance of fixed primary diagnosis models and available comparator representations. (B) Observed performance and 25 fixed-pipeline label-permutation results; these permutations do not repeat the complete nested tuning or historical model search. (C) Representative CD-versus-control calibration curves; threshold-dependent metrics are provided in Source Data. (D) Historical best-of-screen AUC minus pooled nested out-of-fold AUC for the primary model. Outer validation used five stratified folds repeated three times and inner three-fold tuning. Fold-median intervals bootstrap 15 correlated outer-fold summaries and are not independent-cohort CIs. Comparator analyses may use different eligible participants; no matched head-to-head superiority is inferred. No external clinical-classifier transfer is included.

### Supplemental table titles

**Table S1. Clinical and demographic characteristics of the study cohort.** Participant-level demographic and clinical characteristics for the PBMC2026 cohort, reported overall and by non-IBD control, ulcerative colitis, and Crohn’s disease groups. Continuous variables are summarized by median, interquartile range, and range where available; categorical variables are reported as number and percentage.

**Table S2. Milo differential-abundance results for CD versus non-IBD control PBMC neighborhoods.** Full Milo differential-abundance output for the CD versus control comparison across 62,426 PBMC neighborhoods. Columns report log fold change (logFC), average log counts per million (logCPM), F statistic, nominal P value, Benjamini-Hochberg FDR, neighborhood identifier (Nhood), spatial false-discovery-rate-adjusted value (SpatialFDR), neighborhood center cell identifier, assigned immune-cell type, and the fraction of cells in the neighborhood assigned to that cell type. Positive logFC values indicate enrichment in CD relative to control; negative values indicate enrichment in controls.

**Table S3. Milo differential-abundance results for UC versus non-IBD control PBMC neighborhoods.** Full Milo differential-abundance output for the UC versus control comparison across 38,371 PBMC neighborhoods. Columns report logFC, logCPM, F statistic, nominal P value, FDR, Nhood, SpatialFDR, neighborhood center cell identifier, assigned immune-cell type, and cell-type fraction. Positive logFC values indicate enrichment in UC relative to control; negative values indicate enrichment in controls.

**Table S4. Milo differential-abundance results for CD versus UC PBMC neighborhoods.** Full Milo differential-abundance output for the CD versus UC comparison across 66,508 PBMC neighborhoods. Columns report logFC, logCPM, F statistic, nominal P value, FDR, Nhood, SpatialFDR, neighborhood center cell identifier, assigned immune-cell type, and cell-type fraction. Positive logFC values indicate enrichment in CD relative to UC; negative values indicate enrichment in UC.

**Table S5. TCR repertoire model-screen performance across diagnosis, inflammation, and contemporaneous response-status tasks.** The workbook contains 594 model–comparison records and separate summaries for ROC AUC, balanced accuracy, Matthews correlation coefficient, and F1 score. It documents the broader historical screen, including available linear, DeepRC, and diversity/clonality models, rather than replacing the primary nested estimates in Figure 7B or conditional out-of-fold estimates in Figure 7D,F. Evaluation design and eligibility must be read with each record.

**Table S6. BCR repertoire model-screen performance across diagnosis, inflammation, and contemporaneous response-status tasks.** The workbook contains 522 model–comparison records, with task-specific ROC AUC, balanced accuracy, Matthews correlation coefficient, and F1 summaries. It documents historical candidate models and representations, not a uniform nested evaluation of every candidate. The primary diagnosis and conditional clinical-state estimates are reported in Figure 7 and its Source Data; individual records retain their own feature definitions, eligible samples, and evaluation designs.

**Table S7. Core TCR and BCR transcriptional gene signatures.** The workbook contains 17 T-cell and 19 B-cell modules, their identifiers, labels, ordered gene sets, and long-format module–gene associations. Scores use the mean normalized expression of detected signature genes. Focused pathogenic/conventional Th17 and suppressive/reprogramming Treg definitions added in subsequent analyses are specified separately in STAR Methods and the analysis code; Table S7 is not an exhaustive list of every focused analysis-specific variant.

## STAR ★ Methods

### Resource availability

#### Lead contact

Further information and requests for resources and reagents should be directed to and will be fulfilled by the lead contact, John Gubatan.

#### Materials availability

This study did not generate new unique reagents.

#### Data and code availability

At publication, this Resource will include single-cell gene-expression matrices; cell- and participant-level metadata; productive TCR and BCR contig tables; exact-clonotype assignments; paired-TCR neighborhood nodes and edges; BCR SHM and lineage tables; transcriptional-program definitions; participant-level source data; frozen model-input matrices and fold assignments; analysis code; and software-environment files. Sequencing datasets will be deposited in the NIH National Center for Biotechnology Information Gene Expression Omnibus (NCBI GEO) at the time of publication. Analysis code, immuneML specifications, figure workflows, and non-sensitive processed outputs are publicly available at https://github.com/GubatanLab/IBD-Blood-Single-Cell-Immune-Repertoire-Atlas. Version 1.0.0 is permanently archived in Zenodo at https://doi.org/10.5281/zenodo.22866361. De-identified clinical metadata will be shared subject to institutional review board approvals, consent, and data-use restrictions. External datasets used here are available as GSE261334, GSE125527, and GSE301689 [17,28,32].

#### Resource organization and intended reuse

The release will use stable de-identified participant identifiers linked, where available, to cell barcodes, exact-clonotype keys, sequence-neighborhood identifiers, and BCR lineage identifiers. A README, data dictionary, manifest, environment specification, and identifier crosswalk will accompany deposition. Panel-specific source tables and code distinguish analysis units, receptor definitions, eligible denominators, missingness rules, and validation level. Frozen feature matrices, participant-level fold assignments, model specifications, and permutation settings will support benchmarking reuse. Participants without productive receptor recovery are excluded from receptor-dependent analyses rather than assigned zero-valued repertoires.

## Acknowledgments

We wish to thank the patient participants for their engagement and effort to enable this study. J.G. and this project were supported in part by a Doris Duke Physician Scientist Fellowship Award (grant no. 2021091), CZ Biohub Physician Scientist Scholar Award, NIH NIDDK LRP Award (2L30 DK126220), Stanford Translational Research and Applied Medicine (TRAM) Scholar Award, Stanford MCHRI Pediatric IBD and Celiac Disease Research Award, and Mayo Clinic Florida Gastroenterology startup funds.

## Author contributions

Conceptualization, J.G.; Methodology, J.G., J.Y., C.L.-A., and G.K.S.; Formal analysis, J.G., J.Y., J.C., Y.Z., C.L.-A., and G.K.S.; Investigation, R.S.S., J.H., T.F., T.T., P.S., A.H.K., M.G., T.B., O.H.N., S.R., and S.R.S.; Resources, J.G., S.R.S., T.B., O.H.N., M.J.R., and G.K.S.; Data curation, J.G., J.Y., J.H., and R.S.S.; Visualization, J.G.; Writing – original draft, J.G., J.C., Y.Z., Y.H., K.P.; Writing – review and editing, all authors; Supervision, J.G., S.R.S., O.H.N., M.J.R., and G.K.S.; Project administration, J.G., R.S.S., T.B., and O.H.N.; Funding acquisition, J.G.

## Declaration of generative AI and AI-assisted technologies in manuscript preparation

No generative AI was used to create, alter, or analyze primary research data. AI-assisted technology was used to proof read manuscript.

## Key resources table

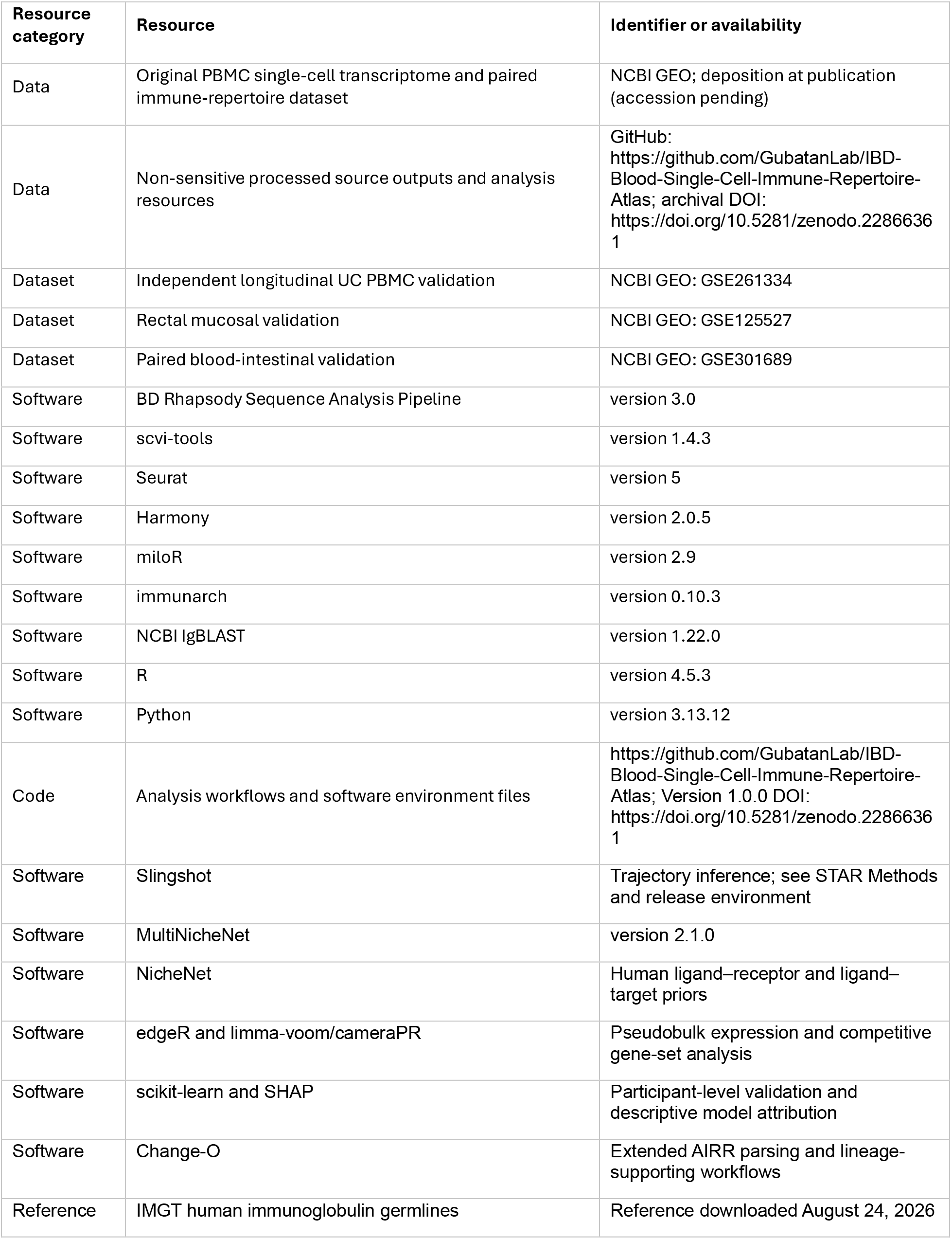

## Experimental model and study participant details

### Human participants

Peripheral blood was obtained from 249 participants: 127 with CD, 87 with UC, and 35 non-IBD controls. Diagnoses were established from clinical, endoscopic, radiographic, and histopathologic criteria. The cohort included 133 female participants (53.4%); median age was 37.0 years (interquartile range [IQR], 28.0-49.0; range, 6.0-79.0). All participants provided informed consent or assent with parent or guardian permission, as appropriate. The study was approved by Stanford University (IRB 28437, 60958, and 52317) and the University of Copenhagen (H-21054506). Clinical variables were linked to sequencing data by de-identified participant identifiers.

### Clinical outcome definitions

Diagnostic analyses compared CD, UC, and non-IBD control status. Objective intestinal inflammation was defined from fecal calprotectin greater than 150 micrograms/g together with endoscopic and/or histologic evidence of inflammation; controls were classified as noninflamed. Biologic response-status analyses included anti-TNF, ustekinumab, and vedolizumab exposure. Response status was assigned at the six-month assessment. Nonresponse required ongoing active symptoms (partial Mayo score greater than 4 or Harvey-Bradshaw Index at least 5) together with fecal calprotectin greater than 150 micrograms/g. Blood used for these analyses was collected at the same six-month assessment; the models therefore classified contemporaneous post-treatment status and were not prospective treatment-response models.

## Method details

### PBMC isolation and single-cell library preparation

Peripheral blood mononuclear cells (PBMCs) were isolated from 5–10 mL of whole blood using Accuspin tubes (Sigma-Aldrich, A2055) layered with Ficoll (GE Healthcare, 14-1440-03). Recovered PBMCs were concentrated in Gibco Recovery Cell Culture Freezing Medium (12648010), cooled at −80 degrees C in controlled-rate freezing containers, and transferred to liquid nitrogen. The BD Human Single-Cell Multiplexing Kit (633781) was used to multiplex up to 12 participant samples per cartridge; cartridges were organized into 10 acquisition series. Approximately 5,000-10,000 cells per participant were loaded onto BD Rhapsody cartridges (633733). Single-cell capture, barcoding, lysis, and cDNA synthesis were performed on the BD Rhapsody Express system. Whole-transcriptome libraries were generated with the BD Rhapsody WTA Amplification Kit (633801), and full-length TCR and BCR libraries with the BD Rhapsody TCR/BCR Amplification Kit (665345). Libraries were sequenced on a NovaSeq X Plus at target depths of 25,000 reads per cell for whole-transcriptome libraries and 5,000 reads per cell for each receptor library.

### Single-cell RNA-seq preprocessing, integration, and annotation

FASTQ files were processed with the BD Rhapsody analysis pipeline on Seven Bridges using default parameters and the GRCh38-PhiX-gencode v29 reference. Filtered UMI matrices were used to fit scVI with scvi-tools version 1.4.3 [36,37], using 50 latent dimensions and 400 training epochs. Technical and sample covariates were supplied where available. The posterior latent representation was imported into Seurat version 5 [38,39] for graph construction, UMAP visualization [40], clustering, and expression analyses. Raw counts were retained; log-normalized values were used for visualization, marker testing, and program scoring. Immune-receptor genes and features dominated by clonotype-specific or technical variation were excluded from variable-feature selection. Residual technical structure was evaluated by participant, acquisition series, sequencing run, site, and diagnosis; Harmony version 2.0.5 [41] was used within lineage-specific workflows when it improved technical mixing without erasing expected immune-state structure. Initial annotations were generated with ScType [42] and published multimodal PBMC references [43,44], followed by manual review of canonical markers. CD4 T cells, CD8 and innate-like T cells, and B cells were subclustered in lineage-specific objects to obtain the level 2 and level 3 states used in clone-aware analyses. Marker-expression matrices used the complete quality-controlled lineage objects. UMAP downsampling reduced overplotting only; balanced trajectory references and analysis-specific eligibility are described separately and must not be equated with the full-cell denominators.

### Differential abundance of PBMC states

Diagnosis-associated abundance was tested with miloR version 2.9 [45]. A k-nearest-neighbor graph was constructed from the 50-dimensional latent representation with k = 20. Partially overlapping neighborhoods were counted by participant, and CD versus control, UC versus control, and CD versus UC contrasts were fitted with negative-binomial models that included acquisition and technical covariates available for the comparison. Multiple testing was controlled with Milo’s graph-based spatial FDR. Neighborhoods with SpatialFDR less than 0.05 were considered differentially abundant. Neighborhoods were assigned to a cell state when the annotation fraction was at least 0.20. Cell-state heatmaps summarized annotation-fraction-weighted neighborhood effects and were descriptive; inference remained at the neighborhood level.

### TCR and BCR processing and clonotype definitions

Receptor reads were grouped by cell barcode, UMI, and chain, error corrected, assembled, and annotated through the BD Rhapsody pipeline [46]. Contigs were assembled with Trinity and annotated using IgBLAST and Bowtie2, with receptor fields represented in AIRR-compliant format [47–50]. Only productive receptors with the chain and sequence fields required for a given analysis were retained. Participants without productive receptor recovery were excluded from receptor-based analyses rather than assigned zero values.

Exact TCR-beta clonotypes were defined within each participant by productive TRBV gene, TRBJ gene, and CDR3 amino-acid identity. Exact paired alpha-beta clonotypes required concordant productive alpha- and beta-chain V genes, J genes, and CDR3 amino-acid sequences; paired gamma-delta and paired heavy-light BCR clonotypes were defined analogously. When more than one contig of a chain type was present, the dominant productive contig selected by the source pipeline was used. Expansion required at least two cells assigned to the same within-participant exact clonotype. General repertoire displays used singleton, small (2-5 cells), medium (6-20), large (21-100), and hyperexpanded (greater than 100) categories; the TCR dose-response analysis used singleton, 2-cell, 3-4-cell, and at least 5-cell categories.

### Repertoire diversity, clonality, gene usage, and isotype

Participant-level repertoire metrics were calculated from positive-count receptor records using immunarch [10,51] and the documented direct-count summaries. Richness was the observed clonotype count; Shannon entropy was H = −Σpᵢ log(pᵢ), and normalized entropy was H/log(S), where S is richness. Clonality was 1 minus normalized entropy in the CDR3-amino-acid immunarch summary. The separately displayed BCR clonal-expansion index applied the same 1 − H/log(S) formula directly to positive-count source rows (set to 1 when S = 1). These are related evenness measures under different source aggregation/filtering, not independent expansion mechanisms. Inverse Simpson diversity, Gini coefficient, and D50 were additional frequency-based summaries. V/D/J usage was count weighted and normalized within participant after allele-suffix removal. Heavy-chain constant-region calls defined isotype; class-switched receptors were assigned to a non-IGHM/non-IGHD class. Participant was the inferential unit for diagnosis comparisons.

### Transcriptional program scoring

Program scores were arithmetic means of log-normalized expression over detected signature genes. Table S7 contains 17 core T-cell and 19 core B-cell signatures, including GZMK inflammatory memory, cytotoxicity, EOMES–ZEB2, naive/central memory, Th1/Tc1, the broader Th17/IL-23 axis, Tph/Tfh help, activated Treg, plasma differentiation, IgA-mucosal, IgG-inflammatory, and antibody-secretion/UPR programs. These signatures overlap and are not mutually exclusive cell identities. Focused conventional-Th17 genes were RORC, CCR6, KLRB1, IL7R, IL23R, CCL20, IL17A, and IL17F; pathogenic-Th17 genes were TBX21, IFNG, CXCR3, GZMK, CSF2, and CCL5. Suppressive-Treg genes were FOXP3, IL2RA, CTLA4, TIGIT, IKZF2, TNFRSF18, LRRC32, and ENTPD1; reprogramming-Treg genes were RORC, KLRB1, CCR6, TBX21, IFNG, CXCR3, GZMK, and CCL5. Internal focused Th17 and Treg scores were summarized within the corresponding annotated populations; external repertoire-wide scores were not equivalent state-restricted tests. Standardization and constant-region exclusion were applied only in the analyses specified below. The focused internal Tph/Tfh-help signature additionally included BATF: CXCL13, PDCD1, ICOS, MAF, IL21, TOX2, TIGIT, CD200, SLAMF6, BATF, CD40LG, CXCR5, BCL6, and SH2D1A. The external core Tph/Tfh signature followed Table S7 and did not include BATF.

### TCR clone-size and program analyses

For each participant and annotated T-cell state, the mean score among singleton cells was subtracted from the score in each expanded clone-size category. Only participant-state strata containing the required comparison were retained; states and CD4/CD8 compartments were then equally weighted so that abundant states did not dominate. Ordered clone-size effects were estimated in participant-fixed-effect models with participant-clustered standard errors. P values were corrected across prespecified programs by the Benjamini-Hochberg procedure.

### CD8 transcriptional trajectory and expansion-by-pseudotime analyses

Conventional-CD8 pseudotime used a consensus graph-geodesic approach in scVI latent space. The reference retained all eligible paired αβ-TCR cells and up to 12 additional cells per participant–state stratum. A 35-neighbor Euclidean graph was locally distance normalized and symmetrized. Root candidates were CD8 naive cells in the upper quartile of the early-memory program; up to 12 participant-distinct roots nearest the candidate latent centroid were retained. Dijkstra distances were rank normalized per root and averaged; remaining transcriptome cells were projected using 15-neighbor distance-weighted averages. Clone position was median pseudotime and span was the 90th minus 10th percentile. Within-participant demeaned regressions related these outcomes to log2 clone size with participant-clustered standard errors, retaining only informative within-participant variation. Displayed participant/bin medians used 3,000 participant-bootstrap CIs. For program profiles, paired cells were partitioned into ten pseudotime quantile bins and scores averaged by participant, bin, and singleton/expanded status. Spline models used four degrees of freedom, participant fixed effects, and participant-clustered errors; joint spline-interaction tests were FDR adjusted across programs. The skeleton is a visualization of transcriptional ordering, not a reconstructed temporal lineage.

### Paired alpha-beta TCR sequence-neighborhood analysis

Participant-specific paired alpha-beta clonotypes were represented by alpha- and beta-chain CDR3 amino-acid 2-mers and 3-mers together with alpha/beta V- and J-gene tokens. Features were constructed independently within each acquisition series. Candidate neighbors were identified by cosine distance; up to five cross-participant neighbors among 40 candidates were retained at distance at most 0.50. Exact paired receptors and within-participant edges were excluded. The baseline analysis contained 43,742 clonotypes and 5,548 cross-participant edges across eight acquisition series; 181 participants contributed at least one edge.

For each transcriptional program, clonotype scores were centered and standardized within participant. Edge concordance was calculated within acquisition series and pooled by random-effects meta-analysis. Uncertainty was estimated by deleting participants and recomputing the graph summary; pooled 95% confidence intervals therefore reflect participant-jackknife variability. Diagnosis-restricted strata required at least 100 edges and six participants. The correlated CD-versus-UC contrast resampled participants within acquisition series 2,000 times. Clone-size-adjusted scores removed a common within-participant linear association with log clonotype size before standardization; singleton analyses excluded expanded clonotypes; dominant-state-adjusted scores removed dominant-state means. Baseline-to-adjusted attenuation was tested by 2,000 participant-block bootstrap replicates while retaining the fixed graph and meta-analysis weights. Paired CDR3-only, alpha-only, and beta-only graphs were analyzed as prespecified sensitivities. Inflammatory-status organization was tested among within-diagnosis IBD edges. Participant inflammatory-status labels were permuted 10,000 times within diagnosis while preserving graph topology and participant clonotype burden. The primary statistic was the observed minus expected fraction of edges joining participants with the same inflammatory status. Participant-delete-one jackknife intervals quantified uncertainty, and edge-type decompositions were corrected across inflamed-inflamed, noninflamed-noninflamed, and mixed-status categories.

### Paired gamma-delta TCR analysis

Paired gamma-delta V-gene architecture was summarized within participant so that participants with many recovered cells did not dominate. Expected V-gene pairing and two-sided P values were obtained by shuffling delta-chain V-gene labels among cells within participant 10,000 times; FDR was controlled across all observed or possible pairs. Program contrasts for TRGV9-TRDV2 versus other paired receptors and expanded versus singleton paired gamma-delta clonotypes were matched within participant and observed state, standardized, and summarized by participant. Confidence intervals used 10,000 participant bootstrap replicates. Two-sided sign-flip tests were exact when feasible and otherwise used 200,000 draws, with FDR correction across the five programs in each contrast.

### BCR expansion, state occupancy, and class switching

BCR state-occupancy analyses in Figure 4B used participant-specific heavy-chain V/J and heavy/light CDR3 amino-acid keys. This paired-chain key does not additionally require light-chain V/J concordance and is distinct from the six-field exact paired definition used for clone-resolved trajectory and lineage-linkage analyses. State fractions were calculated separately among expanded and singleton cells; eligible participants contributed at least five expanded and 20 singleton cells. Median expanded-minus-singleton percentage-point contrasts and 3,000 participant-bootstrap CIs were calculated for ≥2-, ≥3-, and ≥4-cell thresholds; two-sided signed-rank tests were FDR adjusted across states. Heavy-chain isotype occupancy was analyzed separately. Observed switched fractions used rank tests; adjusted switched/unswitched counts used beta-binomial models with diagnosis, age, sex, and log BCR depth and Holm correction across the two disease-control contrasts. Global IgM/IgD/IgA/IgG composition used Dirichlet– multinomial likelihood-ratio tests; centered-log-ratio coefficients describe individual isotype effects. These models measure cross-sectional occupancy, not switching rate or direction.

### State-matched BCR pseudobulk expression

The Figure 4C,D analysis defined expansion by productive heavy-chain V gene, J gene, and CDR3 amino-acid identity within participant, without requiring light-chain recovery. Raw counts were aggregated within participant, level 2 state, and expansion status. For states containing both statuses, counts were scaled to the smaller cell count before aggregation across states; participants required at least five matched cells per status (38 eligible BCR participants). TMM normalization, expression filtering (minimum count 10), and limma-voom models [52,53] used participant fixed effects, expansion status, RNA feature complexity, and mitochondrial fraction, followed by robust empirical-Bayes moderation. Gene-level FDR was controlled globally. Competitive program enrichment used cameraPR [54] on ranked moderated statistics with inter-gene correlation 0.01 and at least three measured genes; FDR was controlled across tested T- and B-cell programs. Figure 4C summarizes median/IQR member-gene log2 fold changes alongside the separate competitive FDR. Figure 4D CIs were approximated as log2 fold change ±1.96 standard errors. This heavy-chain pseudobulk analysis must not be described as an exact paired heavy-light dose-response test.

### B-cell transcriptional trajectory

A balanced reference sampled up to eight B cells per participant and level 3 state (14,868 cells from 249 participants). Slingshot [31] was fitted in the first ten SCVI_50 dimensions, with transitional B cells as the root and atypical-memory, IgM-plasma, IgA-plasma, and IgG-plasma states as terminal constraints (150 approximation points; reweighting enabled). Each branch pseudotime was rescaled to 0–1; the global display used curve-weighted pseudotime, and branch labels used the largest curve weight. IgA-mucosal, plasma-differentiation, and antibody-secretion/UPR scores were standardized across reference cells and summarized within participant and pseudotime bin before participant-bootstrap uncertainty was calculated. The IgA-mucosal trajectory signature retained IGHA1/IGHA2; by contrast, constant-region genes were excluded from the state-residualized lineage-coupling analysis below. Root and endpoint constraints define a transcriptional ordering, not observed temporal differentiation.

### Regional somatic hypermutation and BCR lineage reconstruction

Heavy-chain variable-region sequences were reannotated with NCBI IgBLAST version 1.22.0 against the IMGT human immunoglobulin reference and converted to extended AIRR format with Change-O MakeDb [48,55,56]. Somatic hypermutation (SHM) was calculated from observed-versus-inferred-germline nucleotide mismatches in FWR1, CDR1, FWR2, CDR2, FWR3, and FWR4 over evaluable aligned positions. Participant summaries were calculated for unique heavy sequences, heavy clonotypes with exact paired-light assignments, exact paired heavy-light clonotypes, and paired clonotypes weighted by linked cell abundance. Isotype-stratified analyses were performed separately for IgM-, IgA-, and IgG-assigned heavy sequences. CDR targeting was the within-participant mean SHM in CDR1/CDR2 minus mean SHM in FWR1-FWR4. Full light-chain variable-region alignments were unavailable; paired analyses therefore report clonotype-resolved heavy-chain SHM rather than joint heavy- and light-chain mutation.

Primary heavy-chain lineages were defined within participant by shared V gene, J gene, and junction length, followed by single-linkage clustering at normalized nucleotide Hamming distance ≤0.15; thresholds of 0.10 and 0.20 were sensitivity analyses. Exact light-chain V/J/CDR3-linked subdivisions were annotated and evaluated separately, rather than replacing the primary heavy-chain lineages. Germline-rooted minimum-spanning trees used IMGT-aligned heavy sequences plus the inferred germline, with at most 100 observed nodes per lineage. Germline-inclusive branch length includes germline-to-observed connections; observed-only divergence excludes the germline and requires at least two distinct observed heavy sequences. Isotype and cell state were lineage annotations, not terminal-branch assignments. Co-occupancy of IgA- and IgG-plasma transcriptional states was reported as detected lineage counts, not evidence of an observed class-switch event.

Participant-level disease effects were standardized to the control IQR and compared with rank tests followed by FDR correction. Associations with log-transformed fecal calprotectin were evaluated within CD and UC after rank transformation and residualization for log lineage depth, biologic exposure, and acquisition series. Confidence bands and intervals were obtained by participant bootstrap. These analyses do not infer temporal lineage evolution or antigen specificity.

For lineage-level transcriptional coupling analyses, exact paired heavy-light BCR cells were mapped to germline-aware heavy-chain lineages using participant, heavy-chain V and J genes, and heavy-chain CDR3 amino-acid sequence; ambiguous mappings were excluded. Five prespecified programs were evaluated: IgA-mucosal plasma cell, plasmablast/plasma-cell differentiation, antibody-secretion/unfolded-protein response, IgG-inflammatory plasma cell, and cycling B cell. Immunoglobulin constant-region genes were removed before scoring, scores were centered within participant and annotated B-cell state, and cell scores were averaged within lineage. Models included participant fixed effects and participant-clustered standard errors and adjusted for log lineage abundance, log number of program-scored cells, and lineage mean pseudotime. Continuous outcomes were standardized; class switching was modeled as an absolute probability. Branch-length analyses were restricted to lineages with at least two unique observed sequences, and cross-state occupancy analyses to lineages with at least two mapped cells. Benjamini-Hochberg FDR correction was applied across the 20 program-outcome tests.

### Clone-aware Th17/Treg and helper-state analyses

Th17-like and Treg cells were identified from the lineage-resolved CD4 annotations. Pathogenic and conventional Th17 scores were summarized within Th17-like cells; suppressive and reprogramming Treg scores were summarized within Treg cells; and the Tph/Tfh-help score was summarized across eligible CD4 T cells. Exact paired alpha-beta clonotypes were defined within participant as described above. A mixed clonotype contained at least one Th17-like and one Treg cell. Mixed-clonotype counts were compared with 10,000 within-participant state-label permutations that preserved participant clone sizes and Th17/Treg abundance.

Clone-size associations were estimated among complete CD4 clonotype-state summaries using participant and annotated-state fixed effects, with log2 clonotype size as the predictor and participant-clustered robust standard errors. Expanded-versus-singleton program contrasts were calculated within participant and CD4 state, combined with equal state weights, and summarized with participant-bootstrap 95% confidence intervals and two-sided paired tests. Multiple testing was controlled across the four prespecified Th17/Treg programs within each cohort.

### Cross-compartment T-cell–B-cell coordination

Participant-level T-cell and B-cell repertoire or program features were rank transformed and residualized for prespecified covariates. Global partial Spearman correlations adjusted for diagnosis, age, sex, and receptor depth. Pooled-IBD helper-B-cell analyses adjusted for diagnosis, age, sex, log CD4-cell depth, log B-cell depth, objective inflammation, biologic-response status, and acquisition series. P values were obtained from 5,000 permutations of the B-cell residual within acquisition series, and FDR was controlled across the displayed helper-B-cell tests. Joint pooled-IBD models entered Tph/Tfh help, pathogenic Th17, and suppressive Treg scores simultaneously with diagnosis, age, sex, CD4-cell depth, B-cell depth, objective inflammation, biologic-response status, and acquisition series. Predictors and outcomes were standardized after rank transformation. Heteroskedasticity-robust confidence intervals were used, and FDR was controlled across the 15 helper-by-B-cell terms. Robustness was assessed by omitting each acquisition series in turn.

For the Crohn’s disease cytotoxic-plasma analysis, the combined Tph/Tfh helper score and expanded-TCR cytotoxicity score were entered simultaneously with age, sex, log TCR depth, and log BCR depth. Sensitivity models additionally adjusted for acquisition series, objective inflammation, biologic exposure, and selected cell-state composition. Empirical null intervals were generated by matched-participant re-pairing within acquisition series. In blocked leave-one-series-out validation, each acquisition series was predicted from models fitted to all remaining series. These analyses were cross-sectional and were not interpreted as mediation or direct cellular interaction.

### Independent public-dataset reanalyses

Public accessions, source studies, and processed input files were recorded before reanalysis: longitudinal UC PBMCs, GSE261334 [28]; UC/healthy blood and intestinal samples, GSE125527 [17]; and three CD participants with paired blood/colon CD4 data, GSE301689 [32]. Fixed program definitions were applied to detected genes. For 10x H5 inputs (GSE261334 and GSE301689), scores used log1p counts normalized to 10,000 total gene-expression counts per cell. For the targeted wide-matrix extraction in GSE125527, the denominator was the sum of counts over the union of extracted signature genes rather than the whole transcriptome; those relative scores are not directly comparable with the internal or 10x-normalized scores. Productive contigs were linked by barcode and dominant chains used for exact paired clonotypes within participant and sample; cells without paired receptors were excluded from receptor-specific tests. External analyses used available annotations and receptor-bearing compartments rather than transferring all internal state labels or assuming identical assay sensitivity.

For GSE261334, baseline analyses used ten UC and five healthy participants; longitudinal analyses paired the ten UC participants at baseline and week 6. Exact paired-clone-size bins were singleton, 2 cells, 3–4 cells, and ≥5 cells. For each participant and program, Spearman correlations related ordered bins to mean score; two-sided signed-rank tests compared participant correlations with zero and FDR correction was applied across the external clone-size/program family. Baseline group differences used participant-level rank-sum tests, and changes used paired signed-rank tests. Baseline T– B associations used participant-level Spearman correlations. No internal state matching was imposed on the Figure 2H external test, and no response inference was made without participant-linked response labels. Analyses of GSE125527 compared rectal expansion, exact blood–tissue sharing, and BCR clone-bin/program associations at participant level. GSE301689 blood-shared versus colon-private comparisons were descriptive at n = 3. None constituted external SHM validation or transfer of a frozen clinical classifier. A separate motif analysis evaluated five published GLIPH2-derived IBD specificity patterns [29,30]. Carrier status was compared between IBD and controls in the published European cohort and the internal receptor-evaluable cohort using Fisher’s exact tests and FDR correction. These literature-defined beta-chain motifs were not used to build the paired-chain sequence graph. Their external enrichment tests the published motifs, not the graph’s manuscript-specific neighborhoods or antigen assignments.

### T-cell–B-cell interactome inference

MultiNicheNet version 2.1.0 [33] was applied to raw-count CD4 and B-lineage objects using the human NicheNet ligand–receptor network and ligand–target matrix [34]. The input contained 23,493 genes and 246,480 cells (139,687 CD4 and 106,793 B-lineage cells). Sender/receiver states included Tfh, Th17/Th1–Th17, Treg, naive/transitional B, memory B, atypical-memory B, IgM-plasma, and switched-plasma cells. Pseudobulk differential expression used edgeR [52]; participant–state strata required at least five cells, groups at least four participants, and ligand/receptor prevalence of at least 5% in at least half of samples; the absolute log-fold-change threshold was 0.25. Analysis input inventories contained 116 participants for CD inflammation, 73 for UC inflammation, 151 for CD versus UC within shared acquisition series, and 42 for S6-restricted diagnosis comparisons. These are analysis-level inventories, not uniform denominators for every sender–receiver pair; eligibility depends on state coverage. Acquisition series was included where estimable. Figure 6G displays the inflamed group in each inflammation contrast, CD in CD-versus-UC, and CD or UC in their respective control contrasts. For six selected pathway/sender–receiver categories, the highest-priority eligible interaction was retained per displayed context, yielding 30 summaries across five columns. CD86 rows were restricted to CD86–CD28, Treg TGF-β rows to TGFB1 toward IgM-plasma cells, and Th17 BAFF rows to TNFSF13B. Different contexts can therefore select different B-cell states or receptor genes. Bubble area summarizes priority, not a significance test or calibrated cross-context effect. Exact selections and scores are supplied in Figure 6G Source Data. These analyses infer expression-compatible interactions rather than measured signaling or peptide-specific recognition.

### Immune-repertoire classification and model interpretation

Participant repertoires were encoded as normalized contiguous CDR3 amino-acid or nucleotide 3-mer/4-mer frequencies. The fixed primary diagnosis benchmarks used abundance-weighted amino-acid 4-mers. For the selected models displayed in Figure 7C–G, productive sequences were deduplicated within each participant and specified chain compartment before k-mer counts were normalized; repeated copies of an identical sequence therefore did not add abundance weight in those analyses. Combined-chain representations pooled chain sequences; joint TCR+BCR representations concatenated separately labeled receptor feature blocks. They did not preserve the same-cell chain pairing used in exact-clonotype analyses. The broader screen included immuneML-based and compatible implementations [57]; the validation scripts used scikit-learn logistic regression or linear SVMs [58]. Other repertoire-classification work [59] provides methodological context, not external validation of these models. Participants, not cells or receptor sequences, were assigned to folds, and joint models required both eligible compartments.

Primary diagnosis validation fixed the TCR representation to pooled α+β amino-acid 4-mers and the BCR representation to heavy-chain amino-acid 4-mers, with class-balanced logistic regression. Five stratified outer folds were repeated three times; three-fold inner validation tuned C (0.1, 1, or 10) and chi-squared feature-selection percentile (25% or 100%). Feature dictionaries, filtering, and scaling were fitted to training data. Thus, tuning was nested within fixed representations, not across the complete historical model/representation search. Figure 7B reports the median of 15 outer-fold AUCs with percentile intervals from 4,000 bootstrap resamples of those fold summaries; repeated folds are correlated. Pooled out-of-fold AUC, balanced accuracy at threshold 0.5, Matthews correlation coefficient, F1, and calibration were complementary summaries. Twenty-five label permutations evaluated a fixed pipeline (C = 1; 25% feature selection), not the full nested search; the smallest attainable empirical P was 1/26. Endpoint-specific inflammation and six-month response leaders were selected from the historical screen before five-fold out-of-fold reevaluation; their 95% CIs bootstrap participants. These clinical-state estimates are conditional on selection and do not remove all selection optimism. No external clinical-classifier transfer was completed.

For descriptive interpretation, the selected tabular model was refitted to the complete eligible cohort and SHAP values were calculated [60]. Features were ranked by mean absolute SHAP value. These full-cohort attributions were not used to estimate held-out performance and were not subjected to null-hypothesis testing. Short k-mers were interpreted as distributed model features, not stable clonotypes or antigen-specificity assignments. Joint TCR+BCR analyses were limited to participants with both eligible receptor compartments.

### Quantification and statistical analysis

Analyses were performed in R version 4.5.3 [61] and Python version 3.13.12 [62]. Unless otherwise stated, participants were the independent biological units. Continuous variables are reported as median and IQR, and categorical variables as counts and percentages. Two-group participant-level comparisons used two-sided Wilcoxon rank-sum or paired signed-rank tests as appropriate [63]. Comparisons across three diagnoses used Kruskal-Wallis tests followed by prespecified pairwise tests. Categorical and clonotype-frequency tables used Fisher’s exact tests. Regression covariates, resampling units, and the definition of n are specified in the corresponding subsection and figure legend.

Benjamini–Hochberg correction [64] was applied within the specified testing family unless otherwise stated; FDR <0.05 was considered significant. Participant-level bootstrap, jackknife, and blocked permutation procedures were used for population associations. Exceptions were within-participant receptor/state-label nulls and bootstrap intervals over correlated outer-fold AUC summaries, as specified above. Holm adjustment was used for the two disease-control switching contrasts. Acquisition-series-blocked procedures preserved the stated grouping structure but did not resolve diagnosis–series confounding. Cross-sectional associations, sequence-neighborhood attenuation, pseudotime, and feature attributions were not assigned causal or mediation interpretations.

## REFERENCES

1. Maloy, K.J., and Powrie, F. (2011). Intestinal homeostasis and its breakdown in inflammatory bowel disease. Nature 474, 298–306. 10.1038/nature10208.

2. Khor, B., Gardet, A., and Xavier, R.J. (2011). Genetics and pathogenesis of inflammatory bowel disease. Nature 474, 307–317. 10.1038/nature10209.

3. Graham, D.B., and Xavier, R.J. (2020). Pathway paradigms revealed from the genetics of inflammatory bowel disease. Nature 578, 527–539. 10.1038/s41586-020-2025-2.

4. de Souza, H.S.P., Fiocchi, C., and Iliopoulos, D. (2017). The IBD interactome: an integrated view of aetiology, pathogenesis and therapy. Nat. Rev. Gastroenterol. Hepatol. 14, 739–749. 10.1038/nrgastro.2017.110.

5. Ng, S.C., Shi, H.Y., Hamidi, N., et al. (2017). Worldwide incidence and prevalence of inflammatory bowel disease in the 21st century: a systematic review of population-based studies. Lancet 390, 2769–2778. 10.1016/S0140-6736(17)32448-0.

6. Turner, D., Ricciuto, A., Lewis, A., et al. (2021). STRIDE-II: an update on the Selecting Therapeutic Targets in Inflammatory Bowel Disease (STRIDE) initiative of the International Organization for the Study of IBD (IOIBD): determining therapeutic goals for treat-to-target strategies in IBD. Gastroenterology 160, 1570–1583. 10.1053/j.gastro.2020.12.031.

7. Vermeire, S., Van Assche, G., and Rutgeerts, P. (2004). C-reactive protein as a marker for inflammatory bowel disease. Inflamm. Bowel Dis. 10, 661–665. 10.1097/00054725-200409000-00026.

8. Sands, B.E. (2015). Biomarkers of inflammation in inflammatory bowel disease. Gastroenterology 149, 1275–1285.e2. 10.1053/j.gastro.2015.07.003.

9. D’Haens, G., Ferrante, M., Vermeire, S., et al. (2012). Fecal calprotectin is a surrogate marker for endoscopic lesions in inflammatory bowel disease. Inflamm. Bowel Dis. 18, 2218–2224. 10.1002/ibd.22917.

10. Mhanna, V., Bashour, H., Lê Quý, K., et al. (2024). Adaptive immune receptor repertoire analysis. Nat. Rev. Methods Primers 4, 6. 10.1038/s43586-023-00284-1.

11. Greiff, V., Menzel, U., Miho, E., et al. (2017). Systems analysis reveals high genetic and antigen-driven predetermination of antibody repertoires throughout B cell development. Cell Rep. 19, 1467–1478. 10.1016/j.celrep.2017.04.054.

12. Dash, P., Fiore-Gartland, A.J., Hertz, T., et al. (2017). Quantifiable predictive features define epitope-specific T cell receptor repertoires. Nature 547, 89–93. 10.1038/nature22383.

13. Glanville, J., Huang, H., Nau, A., et al. (2017). Identifying specificity groups in the T cell receptor repertoire. Nature 547, 94–98. 10.1038/nature22976.

14. Stubbington, M.J.T., Lönnberg, T., Proserpio, V., et al. (2016). T cell fate and clonality inference from single-cell transcriptomes. Nat. Methods 13, 329–332. 10.1038/nmeth.3800.

15. Martin, J.C., Chang, C., Boschetti, G., et al. (2019). Single-cell analysis of Crohn’s disease lesions identifies a pathogenic cellular module associated with resistance to anti-TNF therapy. Cell 178, 1493–1508.e20. 10.1016/j.cell.2019.08.008.

16. Smillie, C.S., Biton, M., Ordovas-Montanes, J., et al. (2019). Intra- and inter-cellular rewiring of the human colon during ulcerative colitis. Cell 178, 714–730.e22. 10.1016/j.cell.2019.06.029.

17. Boland, B.S., He, Z., Tsai, M.S., et al. (2020). Heterogeneity and clonal relationships of adaptive immune cells in ulcerative colitis revealed by single-cell analyses. Sci. Immunol. 5, eabb4432. 10.1126/sciimmunol.abb4432.

18. Werner, L., Nunberg, M.Y., Rechavi, E., et al. (2019). Altered T cell receptor beta repertoire patterns in pediatric ulcerative colitis. Clin. Exp. Immunol. 196, 1–11. 10.1111/cei.13247.

19. Wu, J., Pendegraft, A.H., Byrne-Steele, M., et al. (2018). Expanded TCRβ CDR3 clonotypes distinguish Crohn’s disease and ulcerative colitis patients. Mucosal Immunol. 11, 1487–1495. 10.1038/s41385-018-0046-z.

20. Williams, K.G., Kongala, R., Shows, D.M., et al. (2024). T cell repertoire homogeneity and blood-gut overlap in patients with inflammatory bowel disease. Cell. Mol. Gastroenterol. Hepatol. 17, 119–130. 10.1016/j.jcmgh.2023.09.003.

21. Hegazy, A.N., West, N.R., Stubbington, M.J.T., et al. (2017). Circulating and tissue-resident CD4+ T cells with reactivity to intestinal microbiota are abundant in healthy individuals and function is altered during inflammation. Gastroenterology 153, 1320–1337.e16. 10.1053/j.gastro.2017.07.047.

22. Safra, M., Werner, L., Peres, A., et al. (2023). A somatic hypermutation-based machine learning model stratifies individuals with Crohn’s disease and controls. Genome Res. 33, 71–79. 10.1101/gr.276683.122.

23. Pesesky, M., Bharanikumar, R., Le Bourhis, L., et al. (2025). Antigen-driven expansion of public clonal T-cell populations in inflammatory bowel diseases. J. Crohns Colitis 19, jjaf048. 10.1093/ecco-jcc/jjaf048.

24. Kotagiri, P., Rae, W.M., Bergamaschi, L., et al. (2025). Disease-specific B cell clones are shared between patients with Crohn’s disease. Nat. Commun. 16, 3689. 10.1038/s41467-025-58977-y.

25. Martini, G.R., Tikhonova, E., Rosati, E., et al. (2023). Selection of cross-reactive T cells by commensal and food-derived yeasts drives cytotoxic TH1 cell responses in Crohn’s disease. Nat. Med. 29, 2602–2614. 10.1038/s41591-023-02556-5.

26. Palm, N.W., de Zoete, M.R., Cullen, T.W., et al. (2014). Immunoglobulin A coating identifies colitogenic bacteria in inflammatory bowel disease. Cell 158, 1000–1010. 10.1016/j.cell.2014.08.006.

27. Bourgonje, A.R., Andreu-Sánchez, S., Vogl, T., et al. (2023). Phage-display immunoprecipitation sequencing of the antibody epitope repertoire in inflammatory bowel disease reveals distinct antibody signatures. Immunity 56, 1393–1409.e6. 10.1016/j.immuni.2023.04.017.

28. Horn, V., Cancino, C.A., Steinheuer, L.M., et al. (2025). Multimodal profiling of peripheral blood identifies proliferating circulating effector CD4+ T cells as predictors for response to integrin α4β7-blocking therapy in inflammatory bowel disease. Gastroenterology 168, 327–343. 10.1053/j.gastro.2024.09.021.

29. Huang, H., Wang, C., Rubelt, F., et al. (2020). Analyzing the Mycobacterium tuberculosis immune response by T-cell receptor clustering with GLIPH2 and genome-wide antigen screening. Nat. Biotechnol. 38, 1194–1202. 10.1038/s41587-020-0505-4.

30. Chan, J.E., Mohsin, A., Krijgsman, J., et al. (2026). Shared CD4+ T cell receptor specificity groups in Crohn’s disease and ulcerative colitis. JCI Insight 11, e195354. 10.1172/jci.insight.195354.

31. Street, K., Risso, D., Fletcher, R.B., et al. (2018). Slingshot: cell lineage and pseudotime inference for single-cell transcriptomics. BMC Genomics 19, 477. 10.1186/s12864-018-4772-0.

32. Arase, M., Murakami, M., Kihara, T., et al. (2025). Multi-omics uncovers transcriptional programs of gut-resident memory CD4+ T cells in Crohn’s disease. J. Exp. Med. 222, e20242106. 10.1084/jem.20242106.

33. Browaeys, R., Gilis, J., Sang-Aram, C., et al. (2023). MultiNicheNet: a flexible framework for differential cell-cell communication analysis from multi-sample multi-condition single-cell transcriptomics data. Preprint at bioRxiv. 10.1101/2023.06.13.544751.

34. Browaeys, R., Saelens, W., and Saeys, Y. (2020). NicheNet: modeling intercellular communication by linking ligands to target genes. Nat. Methods 17, 159–162. 10.1038/s41592-019-0667-5.

35. Gubatan, J., Sojwal, R.S., Ye, J., et al. (2026). Multi-omics reveal vitamin D regulation of immune-gut microbiome interactions and tolerogenic pathways in inflammatory bowel disease. Cell Rep. Med. 7, 102703. 10.1016/j.xcrm.2026.102703.

36. Lopez, R., Regier, J., Cole, M.B., et al. (2018). Deep generative modeling for single-cell transcriptomics. Nat. Methods 15, 1053–1058. 10.1038/s41592-018-0229-2.

37. Gayoso, A., Lopez, R., Xing, G., et al. (2022). A Python library for probabilistic analysis of single-cell omics data. Nat. Biotechnol. 40, 163–166. 10.1038/s41587-021-01206-w.

38. Stuart, T., Butler, A., Hoffman, P., et al. (2019). Comprehensive integration of single-cell data. Cell 177, 1888–1902.e21. 10.1016/j.cell.2019.05.031.

39. Hao, Y., Stuart, T., Kowalski, M.H., et al. (2024). Dictionary learning for integrative, multimodal and scalable single-cell analysis. Nat. Biotechnol. 42, 293–304. 10.1038/s41587-023-01767-y.

40. McInnes, L., Healy, J., Saul, N., and Großberger, L. (2018). UMAP: Uniform Manifold Approximation and Projection. J. Open Source Softw. 3, 861. 10.21105/joss.00861.

41. Korsunsky, I., Millard, N., Fan, J., et al. (2019). Fast, sensitive and accurate integration of single-cell data with Harmony. Nat. Methods 16, 1289–1296. 10.1038/s41592-019-0619-0.

42. Ianevski, A., Giri, A.K., and Aittokallio, T. (2022). Fully-automated and ultra-fast cell-type identification using specific marker combinations from single-cell transcriptomic data. Nat. Commun. 13, 1246. 10.1038/s41467-022-28803-w.

43. Hao, Y., Hao, S., Andersen-Nissen, E., et al. (2021). Integrated analysis of multimodal single-cell data. Cell 184, 3573–3587.e29. 10.1016/j.cell.2021.04.048.

44. Terekhova, M., Swain, A., Bohacova, P., et al. (2023). Single-cell atlas of healthy human blood unveils age-related loss of NKG2C+GZMB−CD8+ memory T cells and accumulation of type 2 memory T cells. Immunity 56, 2836–2854.e9. 10.1016/j.immuni.2023.10.013.

45. Dann, E., Henderson, N.C., Teichmann, S.A., et al. (2022). Differential abundance testing on single-cell data using k-nearest neighbor graphs. Nat. Biotechnol. 40, 245–253. 10.1038/s41587-021-01033-z.

46. BD Biosciences (n.d.). BD Rhapsody Sequence Analysis Pipeline: TCR and BCR analysis. https://bd-rhapsody-bioinfo-docs.genomics.bd.com/steps/steps_tcr_bcr.html (accessed September 20, 2026).

47. Grabherr, M.G., Haas, B.J., Yassour, M., et al. (2011). Full-length transcriptome assembly from RNA-Seq data without a reference genome. Nat. Biotechnol. 29, 644–652. 10.1038/nbt.1883.

48. Ye, J., Ma, N., Madden, T.L., and Ostell, J.M. (2013). IgBLAST: an immunoglobulin variable domain sequence analysis tool. Nucleic Acids Res. 41, W34–W40. 10.1093/nar/gkt382.

49. Langmead, B., and Salzberg, S.L. (2012). Fast gapped-read alignment with Bowtie 2. Nat. Methods 9, 357–359. 10.1038/nmeth.1923.

50. Vander Heiden, J.A., Marquez, S., Marthandan, N., et al. (2018). AIRR Community standardized representations for annotated immune repertoires. Front. Immunol. 9, 2206. 10.3389/fimmu.2018.02206.

51. Popov, A., Samokhina, M., Balashov, I., et al. (2025). immunomind/immunarch: 0.10.3 [software]. Zenodo. 10.5281/zenodo.17358072.

52. Robinson, M.D., McCarthy, D.J., and Smyth, G.K. (2010). edgeR: a Bioconductor package for differential expression analysis of digital gene expression data. Bioinformatics 26, 139–140. 10.1093/bioinformatics/btp616.

53. Law, C.W., Chen, Y., Shi, W., and Smyth, G.K. (2014). voom: precision weights unlock linear model analysis tools for RNA-seq read counts. Genome Biol. 15, R29. 10.1186/gb-2014-15-2-r29.

54. Wu, D., and Smyth, G.K. (2012). Camera: a competitive gene set test accounting for inter-gene correlation. Nucleic Acids Res. 40, e133. 10.1093/nar/gks461.

55. Lefranc, M.-P., Giudicelli, V., Ginestoux, C., et al. (2009). IMGT®, the international ImMunoGeneTics information system®. Nucleic Acids Res. 37, D1006–D1012. 10.1093/nar/gkn838.

56. Gupta, N.T., Vander Heiden, J.A., Uduman, M., et al. (2015). Change-O: a toolkit for analyzing large-scale B cell immunoglobulin repertoire sequencing data. Bioinformatics 31, 3356–3358. 10.1093/bioinformatics/btv359.

57. Pavlović, M., Scheffer, L., Motwani, K., et al. (2021). The immuneML ecosystem for machine learning analysis of adaptive immune receptor repertoires. Nat. Mach. Intell. 3, 936–944. 10.1038/s42256-021-00413-z.

58. Pedregosa, F., Varoquaux, G., Gramfort, A., et al. (2011). Scikit-learn: machine learning in Python. J. Mach. Learn. Res. 12, 2825–2830. https://www.jmlr.org/papers/v12/pedregosa11a.html.

59. Zaslavsky, M.E., Craig, E., Michuda, J.K., et al. (2025). Disease diagnostics using machine learning of B cell and T cell receptor sequences. Science 387, eadp2407. 10.1126/science.adp2407.

60. Lundberg, S.M., and Lee, S.-I. (2017). A unified approach to interpreting model predictions. Adv. Neural Inf. Process. Syst. 30, 4765–4774. https://papers.neurips.cc/paper_files/paper/2017/hash/8a20a8621978632d76c43dfd28b67767-Abstract.html.

61. R Core Team (2026). R: A language and environment for statistical computing (R Foundation for Statistical Computing). https://www.R-project.org/.

62. Python Software Foundation (2026). Python 3.13.12 documentation. https://docs.python.org/release/3.13.12/.

63. Wilcoxon, F. (1945). Individual comparisons by ranking methods. Biometrics Bull. 1, 80–83. 10.2307/3001968.

64. Benjamini, Y., and Hochberg, Y. (1995). Controlling the false discovery rate: a practical and powerful approach to multiple testing. J. R. Stat. Soc. Series B Methodol. 57, 289–300. 10.1111/j.2517-6161.1995.tb02031.x.

